# Housing laboratory mice deficient for *Nod2* and *Atg16l1* in a natural environment uncovers genetic and environmental contributions to immune variation

**DOI:** 10.1101/741470

**Authors:** Jian-Da Lin, Joseph C. Devlin, Frank Yeung, Caroline McCauley, Jacqueline M. Leung, Ying-Han Chen, Alex Cronkite, Christina Hansen, Charlotte Drake-Dunn, Kelly V. Ruggles, Ken Cadwell, Andrea L. Graham, P’ng Loke

## Abstract

The relative contributions of genetic and environmental factors to variation in immune responses are still poorly understood. Here, we performed a deep phenotypic analysis of immunological parameters of laboratory mice released into an outdoor enclosure, carrying susceptibility genes (*Nod2* and *Atg16l1*) implicated in the development of inflammatory bowel diseases. Variations of immune cell populations were largely driven by environment, whereas cytokine production in response to stimulation was affected more by genetic mutations. Multi-omic models identified transcriptional signatures associated with differences in T cell populations. Subnetworks associated with responses against *Clostridium perfringens, Candida albicans* and *Bacteroides vulgatus* were also coupled with rewilding. Hence, exposing laboratory mice carrying different genetic mutations to a natural environment uncovered important contributors to immune variation.

**One sentence summary:** Natural environment exposure in laboratory mice primarily promotes variation in population frequencies of immune cells, whereas cytokine responses to stimulation are affected more by genetic susceptibility to inflammatory bowel disease.

## INTRODUCTION

Variation in the magnitude of the immune response is an important determinant of susceptibility to pathogen infections, as well as a predisposition to autoimmunity and cancer (Brodin and Davis, 2017; Schirmer et al., 2018). However, immunological studies with laboratory mice typically aim to control variation in order to focus on genetic factors that alter components of both the adaptive and innate immune system. Recent advances in technology and throughput have facilitated a systems immunology approach towards deciphering the relative genetic and environmental contributors to variation of the human immune system (Bakker et al., 2018; Brodin et al., 2015; Li et al., 2016; Schirmer et al., 2016; Ter Horst et al., 2016). By altering the environment in which laboratory mice are conventionally housed, combined with deep phenotypic analysis of immune responses, we describe here a resource dataset that we used to probe the role of genetic mutations versus environmental drivers of heterogeneity in the immune system. Controlled experiments on specific groups with the same genetic mutations exposed to a new environment would be difficult to conduct in human studies.

Recently, there has been a growing effort to characterize wild mice and pet shop mice, which appear to share greater similarity to the immunological state of humans than laboratory mice (Abolins et al., 2017; Beura et al., 2016; Rosshart et al., 2019; Rosshart et al., 2017). We have developed an outdoor enclosure facility that enables us to reintroduce laboratory mice into a more natural environment, a process termed “rewilding” (Leung et al., 2018). The enclosures include soil and vegetation as well as barnlike structures but no other mammals (e.g., no wild conspecifics from whom lab-bred mice might acquire infections). Over the course of several weeks, mice dig burrows and otherwise adapt to their new environment (Supplemental Videos). Previously, we have observed C57BL/6 mice in this environment exhibit a decreased type 2 and increased type 1 response to intestinal nematode (*Trichuris muris*) infection and greater susceptibility to heavier worm burdens (Leung et al., 2018). This facility provides us with a unique opportunity to rapidly alter the housing environment of laboratory mice that carry genetic mutations and be able to recover them for subsequent detailed immunological profiling.

Gene-bacteria associations have been identified in both human and wild mice indicating host genetic determinants of the gut microbial composition across mammals in natural environments (Suzuki et al., 2019). To interrogate the impact of host genetic variants on microbiota and inflammatory bowel disease (IBD) susceptibility in response to environmental changes, we investigated *Atg16l1* and *Nod2* variants which are among the highest risk factors for IBD associated with a dysregulated immune response to gut microbes (Wlodarska et al., 2015; Wong and Cadwell, 2018). *Atg16l1* is a key component of the autophagy pathway, and *Nod2* is involved in bacterial sensing. We and others previously demonstrated that mice harboring mutations in *Atg16l1* or *Nod2* develop signs of inflammation in a manner dependent on exposure to infectious entities (Biswas et al., 2010; Cadwell et al., 2010; Lavoie et al., 2019; Matsuzawa-Ishimoto et al., 2017; Pott et al., 2018; Ramanan et al., 2016; Ramanan et al., 2014). Mice with these particular mutations are sensitive to the presence of murine norovirus (MNV), *Helicobacter* species, and other commensal-like agents that are found in some animal facilities and excluded in others (Biswas et al., 2010; Cadwell et al., 2010; Caruso et al., 2019; Kernbauer et al., 2014; Pott et al., 2018; Ramanan et al., 2014). Since mice deficient in *Atg16l1* or *Nod2* can exhibit dysregulated immune responses to colonization by organisms that are otherwise commensals in wild type animals, we reasoned that they could be an interesting model system to investigate gene-environment interactions in the outdoor enclosure.

We hypothesized that by introducing laboratory mice carrying IBD susceptibility mutations into the outdoor enclosure, the drastic change in environment may trigger alterations in the immune response and the microbiota, which would have greater adverse effects on mutant mice than wildtype mice. Hence, to study the immunological consequences of rewilding, we released *Atg16l1*^*T316A*/*T316A*^, *Atg16l1*^*T316A*/+^, *Nod2*^−/−^ mice in addition to C57BL/6J wild type (WT) mice to determine whether mutations in these genes alter the immune response to microbial exposure in a natural environment. Here we present a detailed systems immunology profile of both rewilded and laboratory mice to further investigate variation in immune response and the relationship with environment and genetic mutations.

## RESULTS

### Study design for immune profiling of rewilded and laboratory mice

From 116 mice released into the outdoor enclosure for 6-7 weeks we recovered 104 mice (25 WT, 28 *Nod2*^−/−^, 27 *Atg16l1*^*T316A*/+^, and 24 *Atg16l1*^*T316A*/*T316A*^) in time for analysis and compared them to 80 matched controls (19 WT, 19 *Nod2*^−/−^, 20 *Atg16l1*^*T316A*/+^, 22 *Atg16l1*^*T316A*/*T316A*^) maintained under specific pathogen free (SPF) conditions (herein referred to as lab mice) (Figure 1A). Blood samples were collected at the time of sacrifice and analyzed by flow cytometry with a lymphocyte panel (Table S1). Additionally, cytokine production in the plasma was assayed (Figure 1A). Fecal samples and cecal contents were collected for microbial profiling, and for reconstitution experiments (Companion study, Yeung *et. al.*). Mesenteric lymph node (MLN) cells were isolated for flow cytometry analysis with the same lymphocyte panel as the blood and an additional myeloid cell panel (STAR METHODS). Immune activity in the MLN was measured through transcriptional profiles by RNA-seq and cytokine production assays in response to microbial stimulations (STAR METHODS). All of the lab mice were sacrificed in one week, whereas the rewilded mice were sacrificed over a 2-week period based on trapping frequency. This collection of multi-parameter datasets from nearly 200 mice provides an opportunity to examine inter-individual variation in innate and adaptive immune cell populations by flow cytometry analysis, evaluate cytokine responses to microbial stimulation, and integrate highly dimensional transcriptomics and microbiota data. In order to perform multi-omic analysis, 81 mice were completely profiled with measurements from all four data types; flow cytometry, MLN RNA-Seq, fecal microbiota and MLN restimulation and cytokine profiles. A systems level analysis also enabled us to quantify effects of environmental influences and IBD susceptibility gene mutations, *Nod2* and *Atg16l1*, on the heterogeneity of immunological parameters among individual mice.

**Figure 1.**
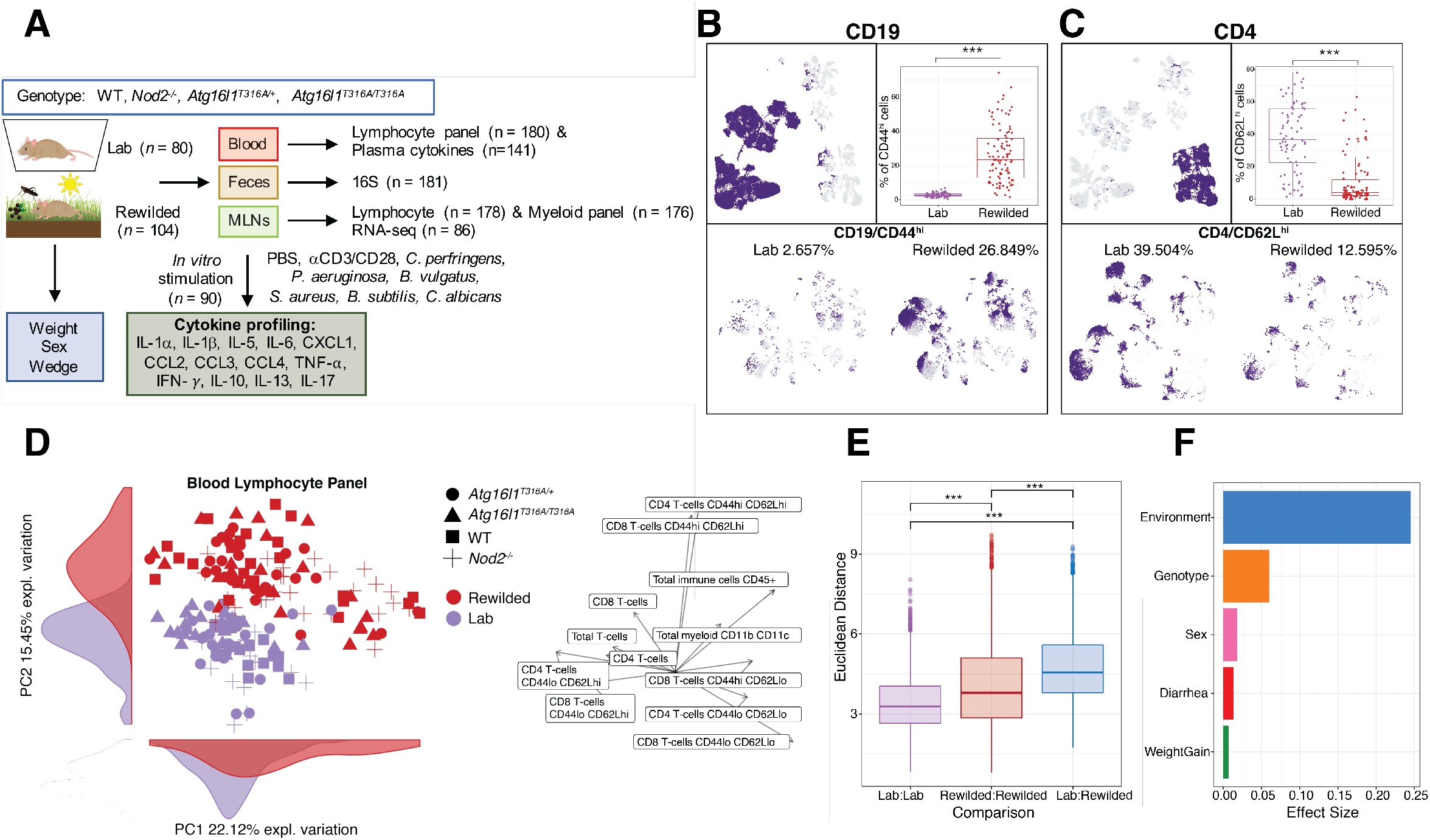
Environmental change drives inter-individual variation in immune cell populations. (**A**) 25 C57BL/6^+/+^, 28 *Nod2*^−/−^, 27 *Atg16l1*^*T316A*/+^, 24 *Atg16l1*^*T316A*/*T316A*^ mice (total=104) were housed in the wild enclosure (Rewilded) for 6-7 weeks and successfully trapped for flow cytometry analysis of lymphoid and myeloid cell populations in the blood and mesenteric lymph nodes (MLNs). Plasma was also collected for cytokine profiles. Age matched 19 C57BL/6^+/+^, 19 *Nod2*^−/−^, 20 *Atg16l1*^*T316A*/+^, 22 *Atg16l1*^*T316A*/*T316A*^ mice (total=80) housed under SPF conditions (Lab) were analyzed as controls. Feces were collected for 16S rRNA sequencing. MLN cells were cultured with indicated stimulates for 48 hours and supernatants were collected for cytokine profiling. Samples that fail quality control are not included in downstream analyses. (**B** and **C**) UMAP projections of ~180,000 CD45+, ~110,000 CD19+, and ~36,000 CD4+ cells on flow cytometry data of the blood from lab and rewilded mice. Events are color-coded according to CD19, CD44 (B), CD4, and CD62L (C) fluorescence level. Box plots show the abundance of CD44^hi^ CD19+ cells (B) and CD62L^hi^ CD4+ cells (C) in the blood of individual lab and rewilded mice. (**D**) Principal component analysis (PCA) of gated immune cell populations in the blood and the density of each population along the principal component (PC). Right panel indicate biplots of the gated immune cell populations that are projected onto PC1 and PC2. (**E**) Box plot of pairwise Euclidean distance measures based on blood immune cell population in the blood of lab versus lab, rewilded versus rewilded and lab versus rewilded mice. (**F**) Bar plot showing the pseudo R^2^ measure of effect size on the entire distance matrix used to calculate the PCA of immune cell populations in the blood (D). \*\*\**P* < 0.001 by Kruskal-Wallis test between groups (B, C) and \*\*\**P* < 0.001 by Mann-Whitney-Wilcoxon test between groups (E).

### Variation of immune cell populations in the blood is largely driven by the environment

In line with previous studies exposing lab mice to feral and pet shop mice (Beura et al., 2016), we observed major differences in CD44^hi^CD62L^lo^ or CD44^hi^CD62L^hi^ T cells (Companion study, Yeung *et. al.*). To visualize the overall landscape of flow cytometry data in an unbiased manner, phenotypic heterogeneity was analyzed on total CD45+ cells per mouse across the blood lymphocyte panel (Table S1) in lab and rewilded mice. We down sampled to 1,000 CD45+ cells from each mouse for nearly 180,000 single CD45+ cell events and single cell flow cytometry data was visualized by uniform manifold approximation and projection (UMAP) (Becht et al., 2019). UMAP analysis was superior to *t*-distributed stochastic neighborhood embedding (t-SNE) (Becht et al., 2019) (Figure S1A) for identifying major immune cell subsets (Figures 1B and 1C; Figures S1B-E). Unexpected clusters of CD44^hi^CD19+ cells were increased in the rewilded mice compared to lab mice (Figure 1B), while lab mice had more CD62L^hi^CD4+ cells (Figure 1C). This unsupervised visualization strategy was useful for identifying unexpected differences (e.g. in the CD19+ compartment and other populations not investigated by (Beura et al., 2016; Rosshart et al., 2019; Rosshart et al., 2017)) in cell clusters of the peripheral blood at a single cell level, which might have been overlooked by traditional flow cytometry gating.

In order to identify known immune cell populations, we employed a traditional gating strategy to systematically quantify the frequencies of immune cell subsets with a lymphocyte panel (Figure S2; Table S2). The blood lymphocyte panel identified 13 lymphocyte populations, a total myeloid cell population (CD11b/CD11c/DX5), and a total CD45+ cell population. From the proportions of these immune cell populations, we examined the inter-individual variation in lab and rewilded mice by principal component analysis (PCA) (Figure 1D). As shown by the separation of lab and rewilded mice along principle component (PC) 1 and 2, there is a strong effect of the environment on lymphocyte populations, and major differences between these populations can be attributed to CD44^hi^CD62L^hi^ CD4 and CD44^hi^CD62L^hi^ CD8 T cells as inferred by examining loading factors along PC1 and PC2 (Figure 1D). The reduced proportion of CD44^lo^ CD62L^hi^ T cells is consistent with some previous reports on wild mice (Abolins et al., 2017; Beura et al., 2016). We also quantified the extent of variability between lab and rewilded mice by measuring the Euclidean distances between mice in our PCA. Notably, the within group distances in lab mice were significantly less than rewilded mice (Figure 1E) suggesting that the natural environment increases variability in immune cell populations. We then evaluated several population factors to identify covariates with the largest effect sizes on immune cell populations (STAR METHODS) (Figure 1F). As expected, the composition and activation state of immune cell populations as determined by cell surface marker expression in the peripheral blood are driven most strongly by environmental differences between the SPF facility and the outdoor enclosure. These differences far outweighed any effects of genetic deficiency in *Nod2* or *Atg16l1* in both environments (Figure 1F). These findings are consistent with the concept that non-heritable influences explain the majority of the variation in the proportion of immune cell subsets as reported in twin studies (Brodin et al., 2015).

### Variation of cytokine production in response to stimulations is driven more by genetic mutations than by the environment

Cytokine production capacity in humans has been reported to be more strongly influenced by genetic than environmental factors (Li et al., 2016). To investigate the relative roles of *Nod2*/*Atg16l1* deficiency and environment on induced responses, we performed *ex vivo* stimulation experiments on total MLN cells from rewilded and lab mice. To examine cytokine responses to microbial and T-cell specific antigens we stimulated with six different microbes (*C. perfringens*, *P. aeruginosa*, *B. vulgatus*, *B. subtilis, S. aureus* and *C. albicans*), αCD3/CD28 beads to activate T cells as well as PBS as a control (Figures 1A and 2A). Despite no significant differences for PBS control samples between lab and rewilded mice (data not shown), we chose to normalize cytokine production per mouse by expressing cytokine data as a fold change levels over PBS (STAR METHODS). A broad array of 13 cytokines was measured: IFN-γ, IL-1α, IL-1ß, IL-13 IL-5, IL-10, IL-17, IL-6, CCL2, CXCL1, CCL3, CCL4 and TNF-α. Average fold changes in cytokine production as compared to PBS controls for both lab and rewilded mice indicate increases in IFN-γ and IL-17 after αCD3/CD28 activation, consistent with the production of these cytokines by T cells in the MLNs (Figure 2A). In general, bacterial antigens induced CCL3, IFN-γ, TNF-α, CCL4, and IL-6 production, while the fungus *C. albicans* induced the regulatory cytokine IL-10 (Figure 2A). Bacterial antigens also induced CXCL1, which was not induced by *C. albicans* or αCD3/CD28 (Figure 2A). As expected αCD3/CD28 was one of the most potent inducers of cytokine production, especially for IFN-γ, CCL4, TNF-α and IL-5, and rewilded mice produced higher levels of these cytokines than lab mice post stimulation (Figure 2B). IL-1α, IL-1β, IL-13 and CCL2 were not appreciably altered by any stimulation (Figure 2A).

**Figure 2.**
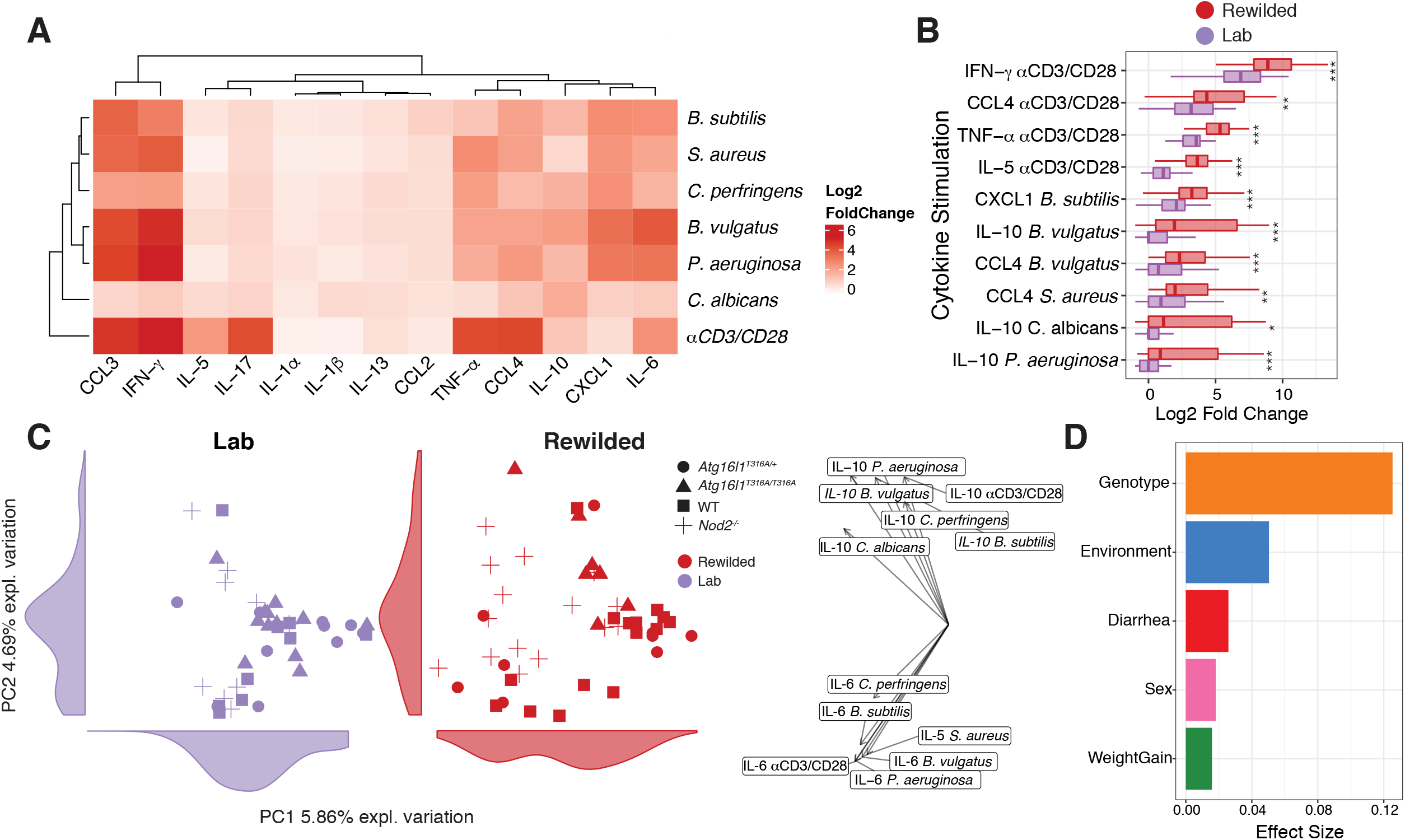
The genetic contribution towards variation in cytokine production in response to microbial stimulation is masked when mice are rewilded. (**A** to **D**) MLN cells from lab and rewilded mice were *ex vivo* cultured with *B. subtilis*, *S. aureus*, *C. perfringens*, *B. vulgatus*, *P. aeruginosa*, *C. albicans*, and αCD3/CD28 beads for 48 hours and the supernatants were assayed for 13 cytokines. (**A**) Heat map of average fold changes compared to cytokine levels of MLN cells cultured with PBS controls from all mice. (**B**) Box plot showing the top 10 log2 fold changes of cytokine responses to stimulation for MLN cells from lab and rewilded mice. (**C**) PCA plot of of fold change cytokine stimulation profiles with histograms of the total variance explained by each PC indicated on the axis. Right panel indicates the loading factor of each stimulated cytokine profile that contributes to each PC. (**D**) Bar plot showing the pseudo R^2^ measure of effect size on the entire distance matrix used to calculate the PCA of cytokine profiles (C). * P <0.05, ** P <0.01, *** *P* < 0.001 by Mann-Whitney-Wilcoxon test between groups (B).

We next used PCA to assess the sources of variation in stimulated cytokine production for both lab and rewilded mice (Figure 2C). Lab and rewilded mice do not clearly segregate along PC1 or PC2 suggesting that environment is not a major source of variation. Effect size measures based on the Euclidean distance indicated genotype as a much stronger source of variation (Figure 2D). The differences in genotype are driven by IL-10 and IL-6 production as indicated by the PCA loading factors (Figure 2C). More specifically, the rewilded *Nod2*^−/−^ mice appear to cluster distinctly from other genotypes, suggesting key differences in cytokine production of rewilded *Nod2*^−/−^ mice (Figure 2C). Genotype also had the greatest effect size on variation in plasma cytokine levels of individual mice (Figure S3).

### Rewilded *Nod2*^−/−^ *mice* exhibit increased cytokine responses compared to wild type rewilded mice

To better understand how *Nod2* and *Atg16l1* deficiency was affecting cytokine production, we performed two-way comparisons and calculated a p-value for each comparison to determine which cytokine producing conditions were most significantly different between WT mice and mice with *Nod2* and *Atg16l1* deficiencies from laboratory and rewilded environments (Figures 3A and 3B). Compared to the lab mice, the rewilded mice had many more significant changes in stimulated cytokine production across *Nod2* and *Atg16l1* deficient mice (Figure 3A and 3B). These changes were largely driven by significant differences in rewilded *Nod2*^−/−^ mice against rewilded WT mice (Figure 3A). Specifically, the *Nod2*^−/−^ mice exhibited increases in IL-17, IL-5 and IL-10 in response to *C. perfringens* suggesting an increased responsiveness to this bacterial stimulant that was only observed in the rewilded condition (Figures 3A and 3C). Production of IL-10 in response to *C. albicans* was also elevated in *Nod2* and *Atg16l1* deficient mice, although *Nod2*−/− did not exhibit greater fungal colonization than WT mice (Figure 3C) (see companion study Yeung. et.al.). Despite these differences in cytokine responses, the rewilded *Nod2*−/− did not shown any signs of increased intestinal inflammation by histology or changes in goblet cell numbers (data not shown), which we have previously shown to be associated with susceptibility to *B. vulgatus* colonization (Ramanan et al., 2016; Ramanan et al., 2014). Together these results indicate that exposure to the outdoor environment per se that induces elevated cytokine responses is not a sufficient hit by itself to trigger intestinal inflammation. Additional insults (i.e. a multi-hit model) may be needed to trigger full blown pathology (Ramanan et al., 2016).

**Figure 3.**
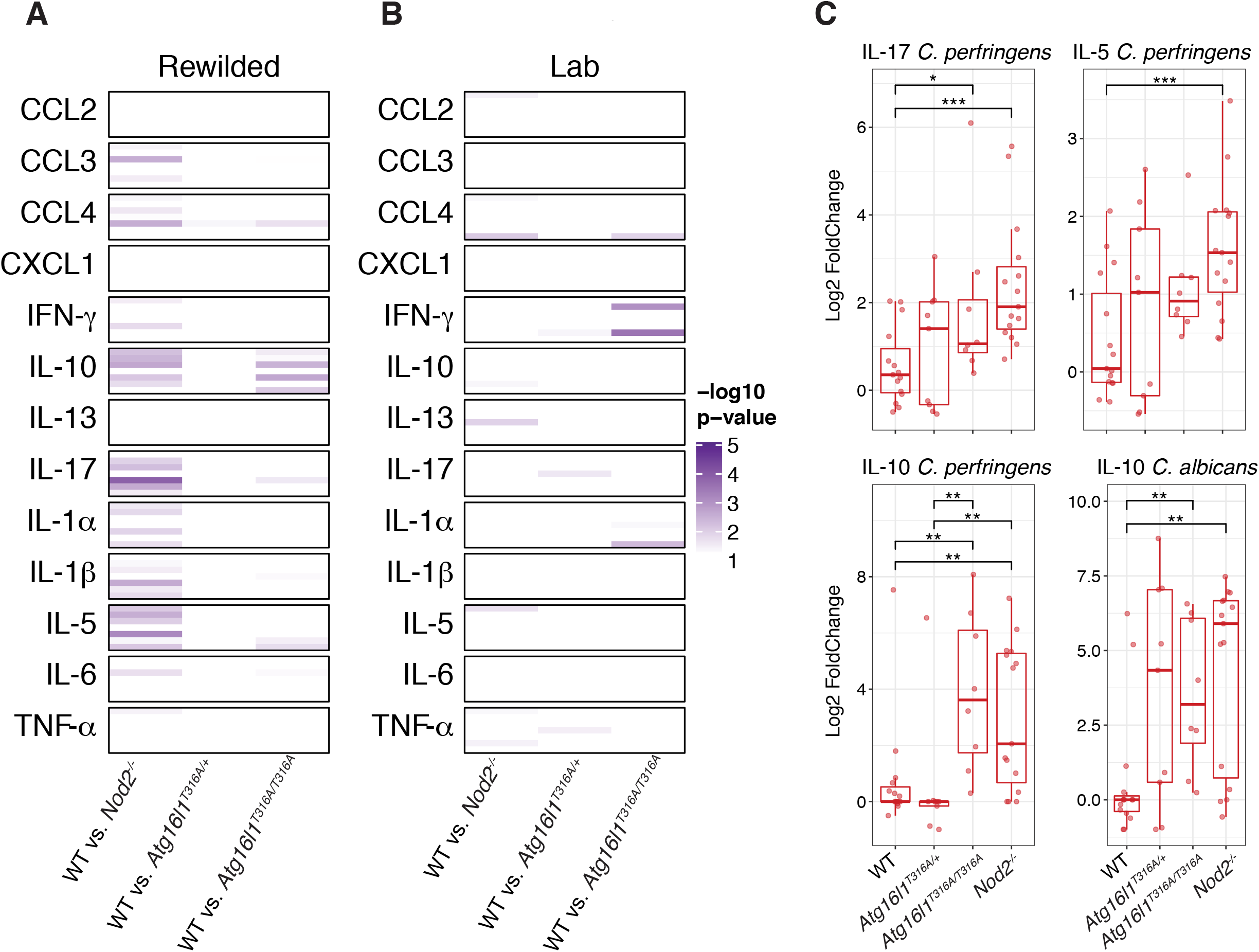
Rewilded *Nod2*^−/−^ mice display greater changes in cytokine production compared to rewilded WT mice. (**A** to **C**) MLN cells from lab and rewilded mice were *ex vivo* cultured with *B. subtilis*, *S. aureus*, *C. perfringens*, *B. vulgatus*, *P. aeruginosa*, *C. albicans*, and αCD3/CD28 beads for 48 hours and the supernatants were assayed for 13 cytokines. (**A** and **B**) Heatmap of significant p-values from Mann-Whitney-Wilcoxon test between group results comparing fold changes of stimulated cytokine production in rewilded (**A**) and lab (**B**) WT mice against *Nod2*^−/−^, *Atg16l1*^*T316A*/+^ and *Atg16l1*^*T316A*/*T316A*^ mice respectively. Within each cytokine condition the order of stimulation remains constant as follows; *B. subtilis*, *B. vulgatus. C. albicans, αCD3/CD28, C. perfringens*, *P. aeruginosa*, *S. aureus*. (**C**) Representative box plots of the most significant changes between WT and *Nod2*^−/−^ mice. * *P* <0.05, ** *P* <0.01, *** *P* < 0.001 by Mann-Whitney-Wilcoxon test between groups (C).

### An integrated classification model identifies features predictive of environment and genotype-specific effects

In addition to stimulated cytokine profiles, MLN cells were assessed for immune cell populations by flow cytometry analysis (Figure S4 and Table S2) and gene expression by RNA-seq analysis (Figures S5A and S5B; Table S2). Consistent with the peripheral blood, immune cell frequencies in MLNs showed a strong effect of environment (Figure S6). Interestingly, a group of rewilded mice all sacrificed at week 7 of rewilding clustered separately in the PCA analysis of lymphocyte populations (Figure S6B). Between week 6 and 7 of rewilding it is possible specific events in the enclosure (e.g., a severe thunderstorm, high humidity and fungal blooms observed during that week), could have resulted in changes in T cell activation. Similar to blood lymphocyte populations, RNA-seq analysis (Figures S5A and S5B) and 16S rRNA profiling (Figures S5C-E; see companion study by *Yeung et al*.) showed some environmental effects on variations. This large amount of data prompted us to use a multi-omic classification model that could prioritize features predictive of environment and genotype-specific effects, as well as identify relationships between these features.

We integrated multi-omic data from flow cytometry populations, gene expression, cytokine, and microbial profiles (Table S2) in a subset of mice (n-=81) which had measures for all four data types. A concatenated matrix (Table S3) of these features was used to build a random forest classification model (STAR METHODS) to distinguish samples by environment or genotype (Figure 4). The contribution of each feature to the model was also assessed to identify features predictive of environment or genotype-specific effects (Figures 4A and 4C). When classifying the environment, a hub of transcription factors (including *Zfp36*, *Atf4*, *Fos*, several *Jun* and *Klf* family members) and *Cd69* are strong predictive features of environment (Figure 4A). Pairwise correlation analysis between these top features also reveals associations to CD44^hi^CD62L^hi^ populations for CD4+ and CD8+ T cells in the blood and MLN, as well as IL-5 and TNF-α production in response to anti CD3/CD28 stimulation (Figure 4B). For the genotype model, our classification was not as accurate as the environment, as measured by area under the receiver operator curve (AUC) of 0.78 versus 1 on the test set (STAR METHODS). Pairwise correlation analysis revealed fewer associations between these features and overall less connected nodes (Figure 4D). Instead of cytokines associated with T cell activation by αCD3/CD28, IL-17 responses to microbial antigen (*B. vulgatus* and *P. aeruginosa*) stimulation were correlated (Figures 4B and 4D). Interestingly, expressions of a number of ribosomal proteins (*Rps* and *Rpl* proteins) were also predictive of genotype (Figures 4C and 4D). In summary, we identified a network of transcription factor associated with T cells in the MLNs based on transcriptional profiling data that is strongly predictive of the effects of the different environments.

**Figure 4.**
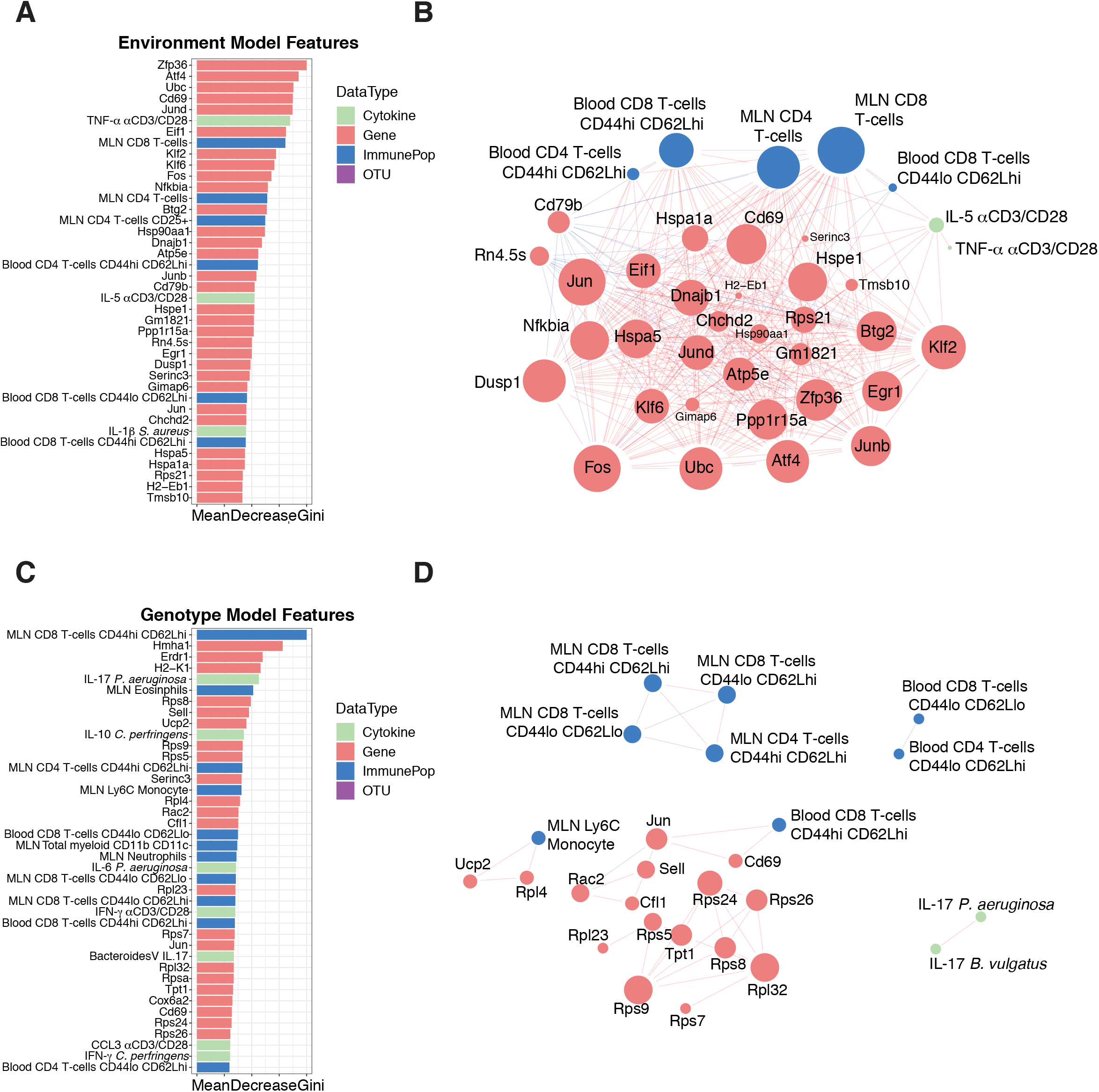
Multi-omic classification models identify predictive features of environment and genotype. (**A**) Bar plot of the top 40 features predictive of the environment and (**B**) interaction network analysis of correlations (R^2^ > 0.6 or R^2^ < −0.6) between features (genes, immune cell populations, OTUs or cytokines) that are predictive of environmental changes. (**C**) Bar plot of the top 40 features predictive of the genotype and (**D**) network analysis of the associations between features that are associated with genotype differences. Data types are colored according to cytokine profiles (green), genes (red), immune populations (blue), and OTUs (purple). The size of the nodes in B and D are scaled proportionally to the number of connections with other nodes in each network.

### Integrative network analysis identifies co-regulated modules in rewilded mice

To evaluate the interactions between data types, we next utilized an unsupervised sparse partial least squares (sPLS) regression model to integrate the different data types into a multi-component network (Figures 5A and S7). To condense the size of our network we collapsed our gene expression profiles according to known Gene Ontologies (STAR METHODS) (2015; Ashburner et al., 2000) for immune system processes in *Mus musculus*, which also focuses our attention on the most relevant genes. In short, we compared our gene expression profiles to all child terms of the Gene Ontology term “immune system process” (GO:0002376) and collapsed our genes into modules based on these pre-defined ontologies. 954 genes were used to generate 91 specific gene ontologies from our gene expression data all related to the parent gene ontology “immune system process”. These modules were used as inputs alongside the cytokine profiles, microbial profiles and flow cytometry populations to generate an unsupervised multi-omic network. sPLS-regression models were built pairwise between each data type to generate a 188-node covariance network with 577 total connections between the four different types of features (Figures 5A and S7). In the resulting model, all microbial taxa fell below our 0.6 covariance threshold and were therefore removed (Table S4). This indicated that bacterial composition, determined by 16S sequencing alone, was not an important component of the network. Perhaps other strategies to assess microbial function (e.g. transcriptomics, metagenomics or metabolomics) would have yielded more connections to immune function.

**Figure 5.**
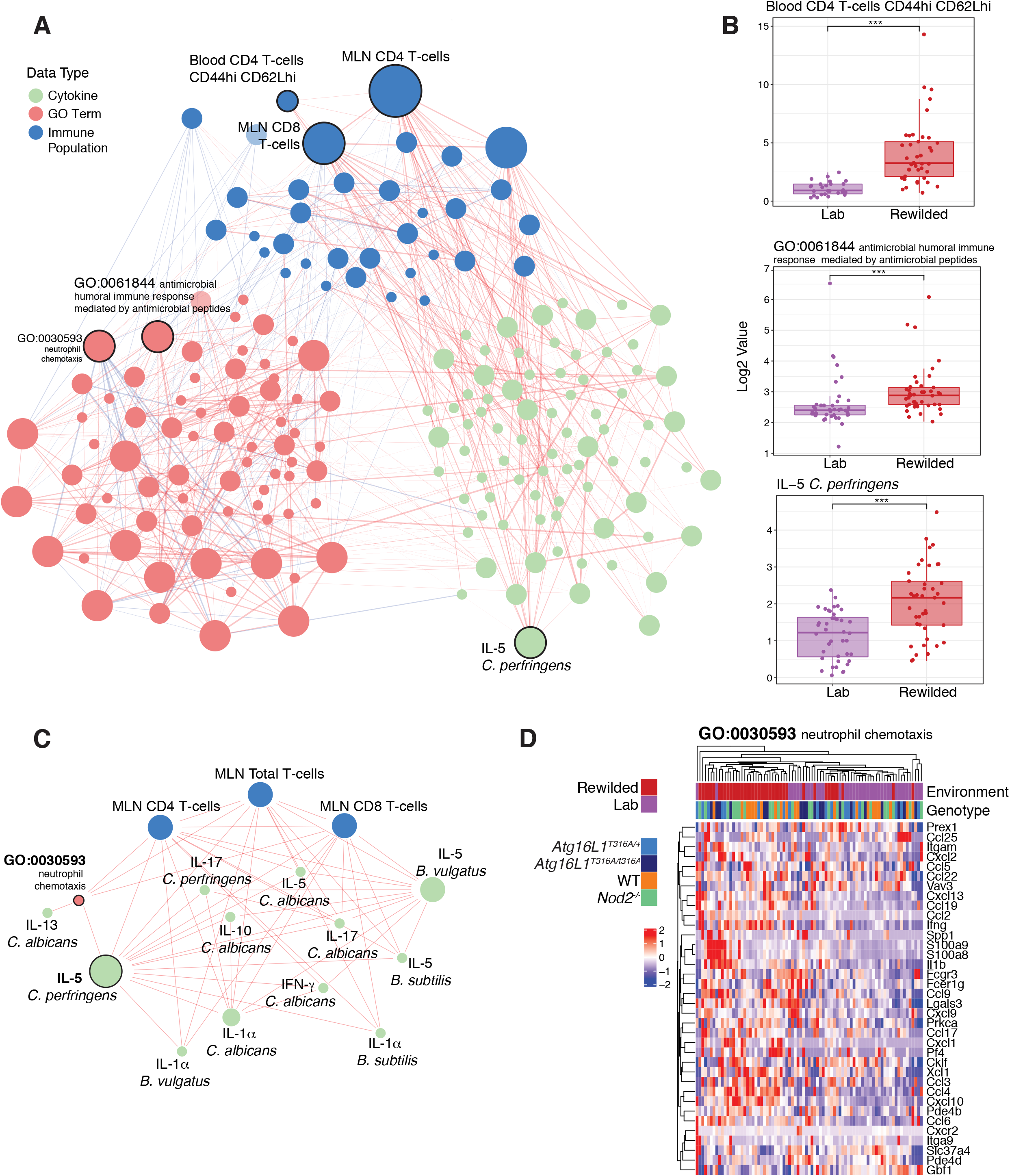
An unsupervised multi-component model identifies a *C. albicans* and *C. perfringens* response network associated with environmental changes. (**A**) Data matrices from flow cytometry data, transcriptional profiles, microbial profiles and stimulated cytokine profiles were used as inputs for a sPLS-regression model to generate a covariance network. (**B**) The most connected nodes, Blood CD4 T-cells CD44hi CD62Lhi, IL-5 in response to *C. perfringens*, and GO:0061844 (antimicrobial humoral immune response mediated by antimicrobial peptides) were evaluated for a difference between lab and rewilded mice populations. (**C**) The IL-5 *C. perfringens* subnetwork was derived from the network and re-calculated to highlight the connections most related to IL-5 production. (D) GO: 0030593 (neutrophil chemotaxis) was identified as a significant connection in the subnetwork and the genes found in that GO term were plotted as a heatmap. Color bars indicate environment and genotype. The size of the nodes in (**A**) and (**C**) are scaled proportionally to the number of connections with other nodes in each network. Data types are colored according to cytokine profiles (green), GO term (red), immune populations (blue). * P <0.05, ** P <0.01, *** *P* <0.001 by Mann-Whitney-Wilcoxon test between groups (B).

While total CD4+ and CD8+ T cell populations in the MLN were among the most connected nodes of the network specific gene ontologies, cytokine responses and other immune populations were also strongly interconnected (Figure 5A). As expected from earlier analysis (Figure 1), the blood CD44^hi^ CD62L^hi^ CD4 T cell population was strongly interconnected and enriched in rewilded mice (Figures 5A and 5B). Additionally, by collapsing our gene expression profiles into annotated gene ontologies, we identified a module (GO:0061844: antimicrobial humoral immune response mediated by antimicrobial peptides) as the most connected gene module that was enriched in rewilded mice (Figures 5A and 5B). Interestingly, IL-5 production in response to *C. perfrigens* stimulation was the most connected cytokine response and was found to be higher in rewilded mice (Figures 5A and 5B; Figure S7). This node is part of a 15-node subnetwork connected to total CD4 and CD8 T cell populations in the MLN, several other cytokine responses and the GO:0030593: neutrophil chemotaxis module (Figure 5C). This module is of particular interest because of the increased neutrophilia observed in the rewilded mice (Yeung et.al. companion paper), which is driven by increased fungal colonization. Indeed, this module is tightly linked with cytokine responses to *C. albicans* antigen stimulation (Figure 5C). While expressions of genes in this module are increased by the rewilding environment, there does not appear to be genotype specific differences regulating expression of the genes in this module (Figure 5D). This multi-omics network analysis therefore substantiated the omics-by-omics inferences (Figures 1 and 2) but also provided new insight by identifying an important role for *C. albicans* responses during rewilding, which is more fully addressed in the companion study (Yeung et.al. companion paper).

## DISCUSSION

In summary, we found that changing the environment profoundly contributes to the inter-individual variations on immune cell frequencies, whereas cytokine responses to pathogen stimulation were more affected by genetic deficiencies in the IBD susceptibility genes *Nod2* and *Atg16l1*. These observations are consistent with human twin studies whereby the majority of cell population influences (measured by flow and mass cytometry) are determined by non-heritable factors (Brodin et al., 2015), whereas cytokine production capacity in response to stimulation may be more strongly influenced by genetic factors (Li et al., 2016). Another large human study indicated that features of innate immunity were more strongly controlled by genetic variation than lymphocytes, which were driven by environmental effects (Patin et al., 2018). Thus, although studies of lab mice, by design, often emphasize genetic effects on immune phenotype (including cell population distributions), here we show that even brief exposure to a natural environment renders predictors of immune phenotype in mice a better match for predictors of human immune phenotypes. Additionally, our dataset described here will be a useful resource for other investigators to delve into the immunological consequences of rewilding.

An attractive hypothesis arising from these findings is that perhaps environment is a primary driver of the composition of the immune system, but genetics is a stronger driver of per-cell responsiveness. However, in this study we only investigated how *Nod2* and *Atg16l1* influenced the immunological parameters we measured. To test this hypothesis further would require us to release mice from diverse genetic backgrounds into the outdoor enclosure in a similar experiment. This experiment only examined the effects of *Nod2* and *Atg16l1* deficiency in the C57BL/6 background. For example, analysis of macrophage activation from five different strains of mice that provided genetic variation on the order of the human population yielded considerable insights into how genetic variation affects transcriptional regulation mechanisms (Link et al., 2018). Alternatively, mice from the collaborative cross may enable high-resolution genome mapping for such complex traits as cytokine production in response to stimulation (Noll et al., 2019). Studies of wild animal populations may also determine if per cell cytokine responses show a higher genetic variance than do immune cell population distributions. Hence, the combination of new environmental challenge strategies such as rewilding of mouse models, combined with modern multi-omic approaches in fully wild systems, should enable us to better define determinants of immune variation at a molecular level.

This study was initially designed to test the hypothesis that *Nod2* and *Atg16l1* deficient mice may respond to the re-wilding environment in an adverse way. Our previous studies had found that *B. vulgatus* colonization of *Nod2* and norovirus colonization of *Atg16l1* deficient mice predisposed them to intestinal inflammation. Hence, we wanted to examine if these mice with mutations in genes associated with the development of IBD would be associated with environmental triggering of intestinal inflammation. However, we did not observe significant differences in intestinal inflammation based on histology for either the *Nod2* or *Atg16l1* deficient mice, although we find an elevated production of cytokines in the *Nod2* deficient mice in response to the rewilding environment. While there were not significant differences in intestinal inflammation, it is possible that additional insults (i.e. a multi-hit model) are needed to trigger intestinal inflammation that is sufficiently severe to be discernable by histologic examination. In our previous studies, an additional piroxicam (for the *Nod2* deficient mice (Ramanan et al., 2016) or dextran sodium sulfate (DSS) (for the *Atg16l1* deficient mice (Matsuzawa-Ishimoto et al., 2017)) insult was required to drive pathogenesis of the *B. vulgatus*/norovirus colonized mice.

A systems level analysis identified interconnected networks of transcriptional signatures, immune cell populations and cytokine profiles to microbial stimulation, highlighting a potentially important role for fungal stimulation from rewilding (Companion study, Yeung *et. al.*). This unanticipated result may have been missed through conventional comparisons. Increased colonization by commensal fungi is perhaps the most striking environmental effect of the rewilding experiment that we performed. While *Nod2* deficiency did not affect fungal colonization, *Atg16l1* deficient mice are more susceptible (Companion study, Yeung *et. al.*). Future studies with mice from more genetically diverse backgrounds will better define the genetic basis of host susceptibility to commensal fungi colonization through rewilding and determine how this may affect the innate and adaptive immune responses.

Hence, rewilding laboratory mice of different genetic backgrounds and careful monitoring of other non-heritable influences (e.g. differential behavior via which divergent immunological experience of individuals may accrue, by analogy with causes of divergence in neurological phenotype (Freund et al., 2013) observed during rewilding (Cope et al., 2019) could be a powerful new approach towards dissecting drivers of immune response heterogeneity under homeostatic as well as during disease settings or infectious challenges.

## Supporting information

TableS1 and STAR METHODS

## Acknowledgements

We wish to thank William Craigens, Daniel Navarrete Prado, Allison Lee, and Veena Chittamuri for assistance with trapping and husbandry in the field, the PU Lab Animal Resources staff for logistical support. We wish to thank the NYU School of Medicine Flow Cytometry and Cell Sorting, Microscopy, Genome Technology, and Histology Cores for use of their instruments and technical assistance (supported in part by National Institute of Health (NIH) grant P31CA016087, S10OD01058, and and S10OD018338).

## Author contributions

Design of experiments, data analysis, data discussion, and interpretation: J.D.L., J.C.D., F.Y., K.C., A.L.G, and P.L.; primary responsibility for execution of experiments: J.D.L., F.Y., C.M., J.M.L., Y.H.C., A.C., C.H., and C.D.D.; MLN cell RNA and 16S analysis: J.C.D., K.V.R., J.D.L., and F.Y. Supervised and Unsupervised machine learning model analysis: J.C.D., K.V.R., and J.D.L. All authors discussed data and commented on the manuscript.

## Data and materials availability

Raw sequence data from 16*S*, ITS, and RNA sequencing experiments are deposited in the NCBI Sequence Read Archive under BioProject accession number PRJNA559026 and gene expression omnibus (GEO) accession number GSE135472.

---

**Figure S1:**
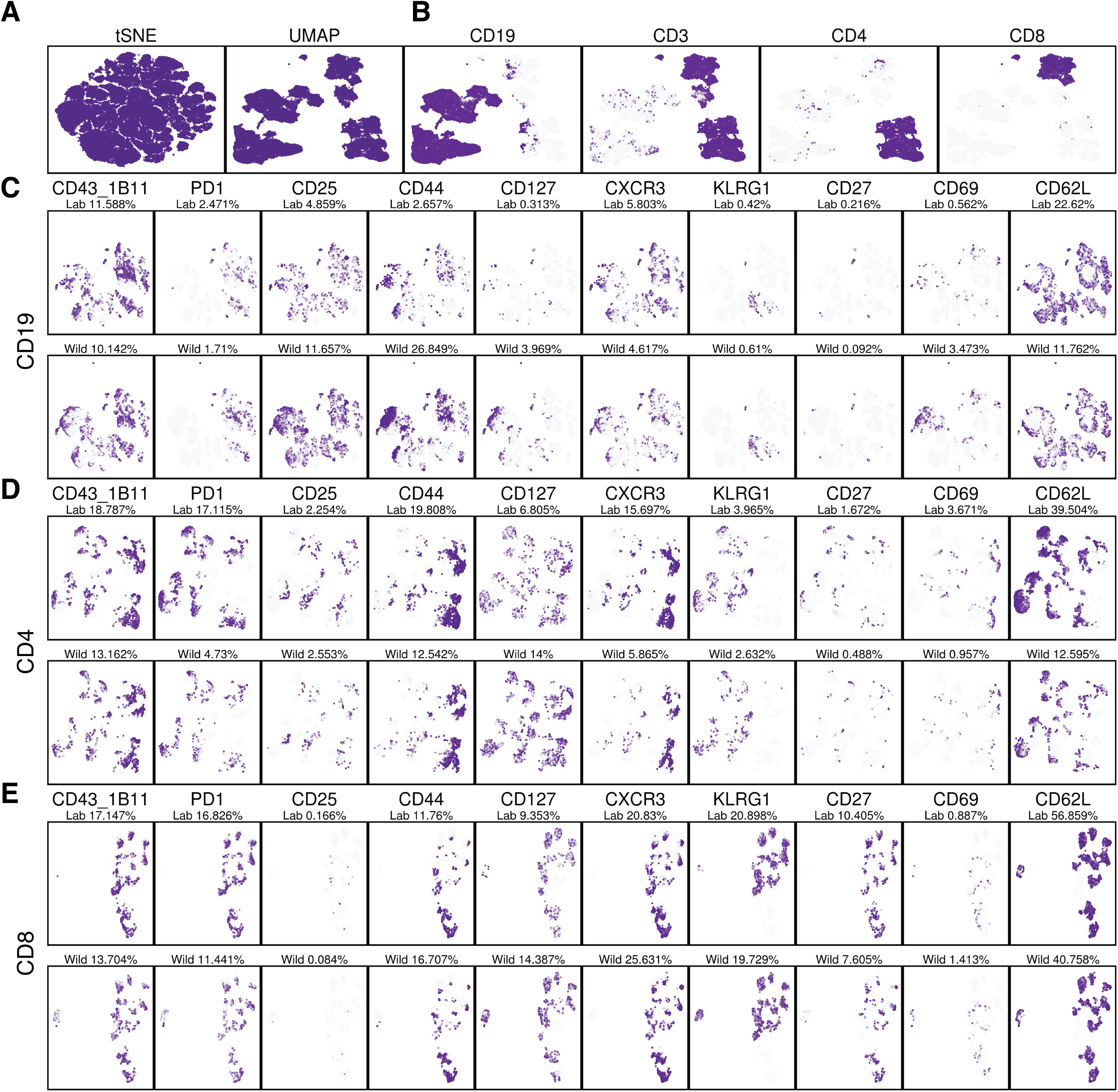
Identification of immune cell populations that differ between lab and rewilded mice with UMAP. (**A**) Projection of ~180,000 CD45+ cells visualized by tSNE vs UMAP. (**B**) Identification of the major lymphocyte populations on UMAP based on CD19, CD3, CD4 and CD8 expression. (**C** to **E**) Expression of cell surface molecules on CD19+ cells (C), CD4+ cells (D) and CD8+ cells (E). In (C), (D) and (E), expression of 1B11, PD1, CD25, CD44, CD127, CXCR3, KLRG1, CD27, CD69, and CD62L are shown as high (purple) or low (grey) from cut-off values based on overall distribution of fluorescence.

**Figure S2:**
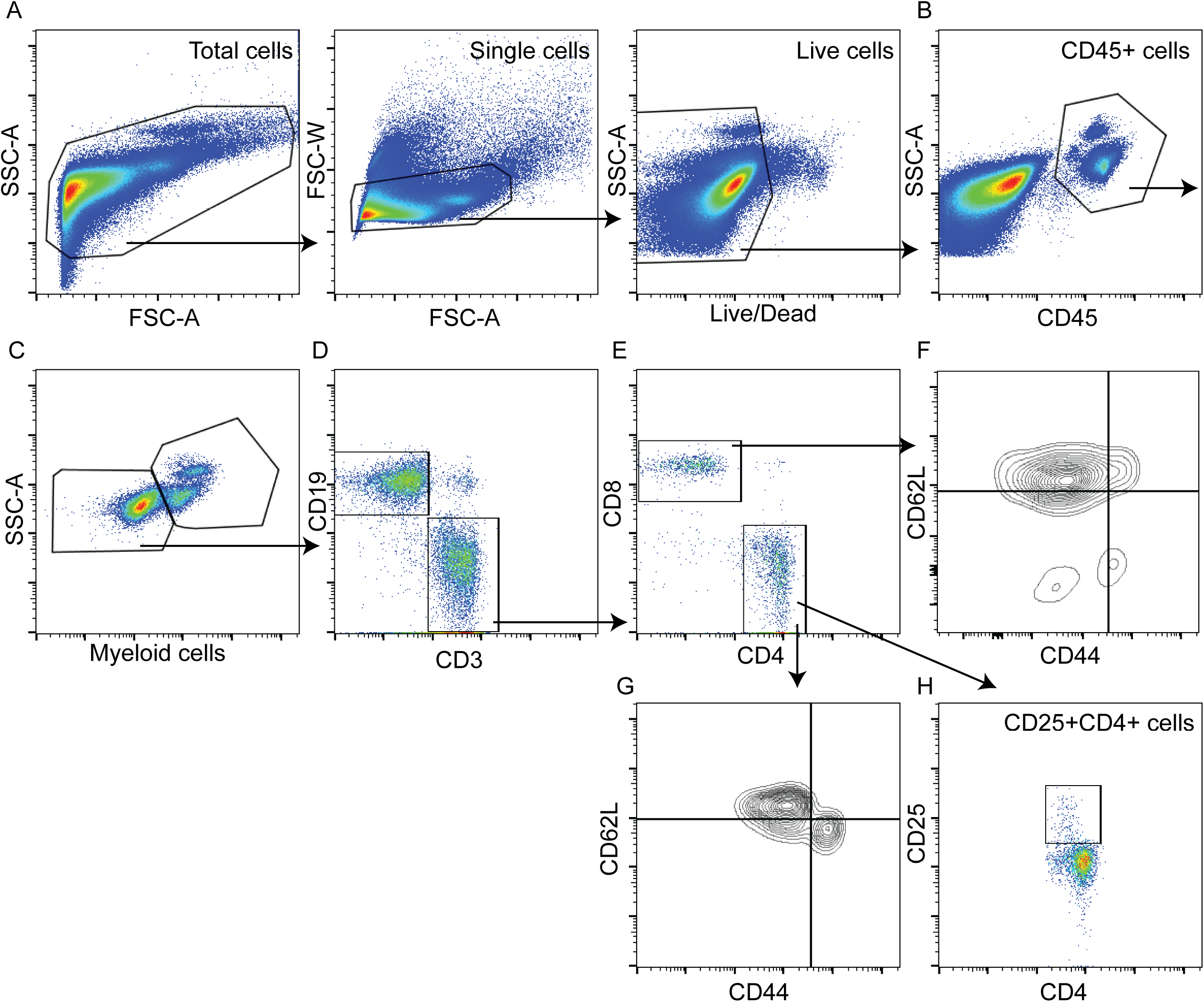
Flow cytometry gating strategy for the quantification of lymphoid populations in the blood of lab and rewilded mice. (**A** to **H**) Representative flow cytometry plots illustrating gating strategy (A) of lymphoid populations in the blood including the total CD45+ cells (B), total myeloid cells (C), total B and T cells (D), total CD4 and CD8 T cells (E), CD44^lo^CD62L^hi^, CD44^hi^CD62L^hi^, CD44^hi^CD62L^lo^, CD44^lo^CD62L^lo^ CD8 T cells (F), CD44^lo^CD62L^hi^, CD44^hi^CD62L^hi^, CD44^hi^CD62L^lo^, CD44^lo^CD62L^lo^ CD4 T cells (G), and CD25+CD4+ T cells (H).

**Figure S3:**
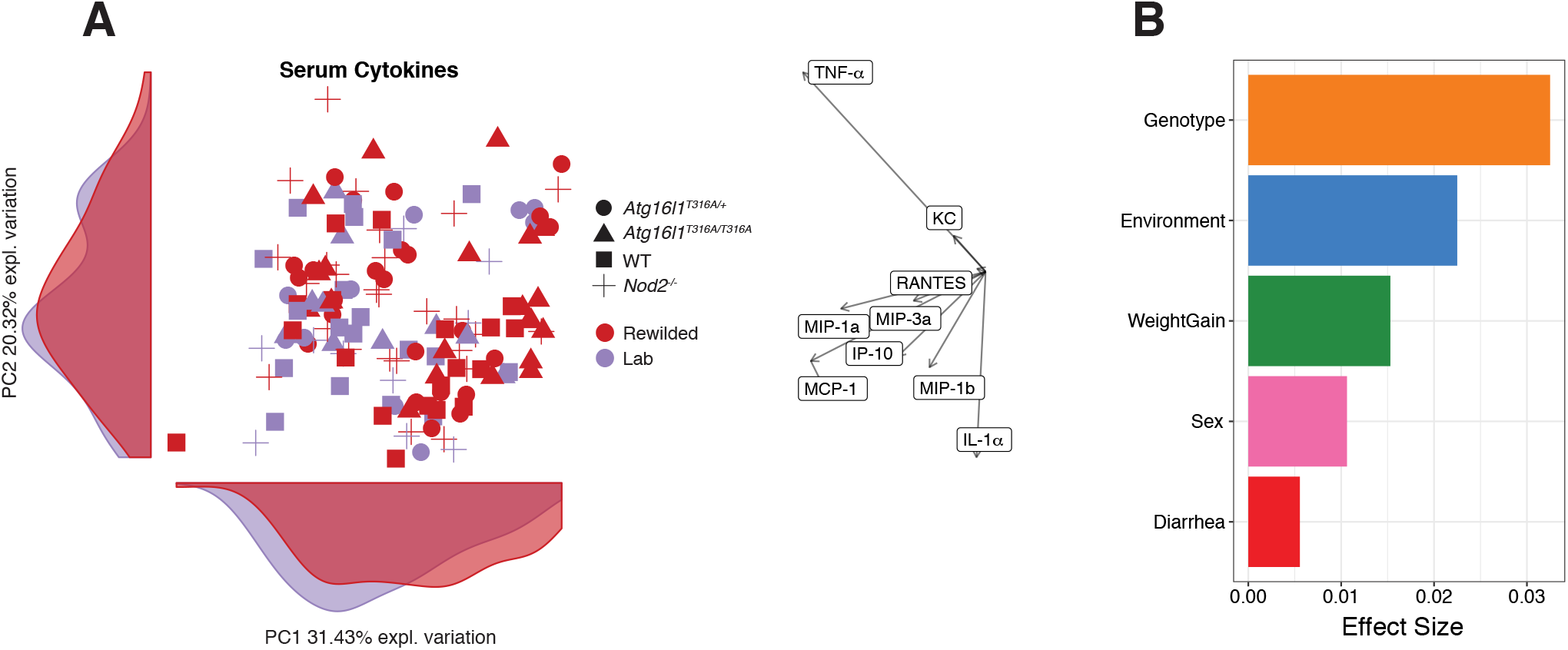
A genetic contribution towards variation in plasma cytokine levels of lab and rewilded mice. (**A** and **B**) Levels of RANTES, MIP-3a, KC, TNF-α, MCP-1, IP-10, MIP-1a, MIP-1b, IL-1α, IL-6 cytokines were measured from the plasma of lab and rewilded mice at the time of sacrifice with bead-based immunoassays (LEGENDplex). (**A**) PCA of plasma cytokine levels in the blood of individual mice and the density of each population along the principal components (PC). Right panel indicate biplots of the plasma cytokines are projected onto PC1 and PC2. (**B**) Bar plot showing the pseudo R^2^ measure of the effect size of different variables on the variance of plasma cytokine levels.

**Figure S4:**
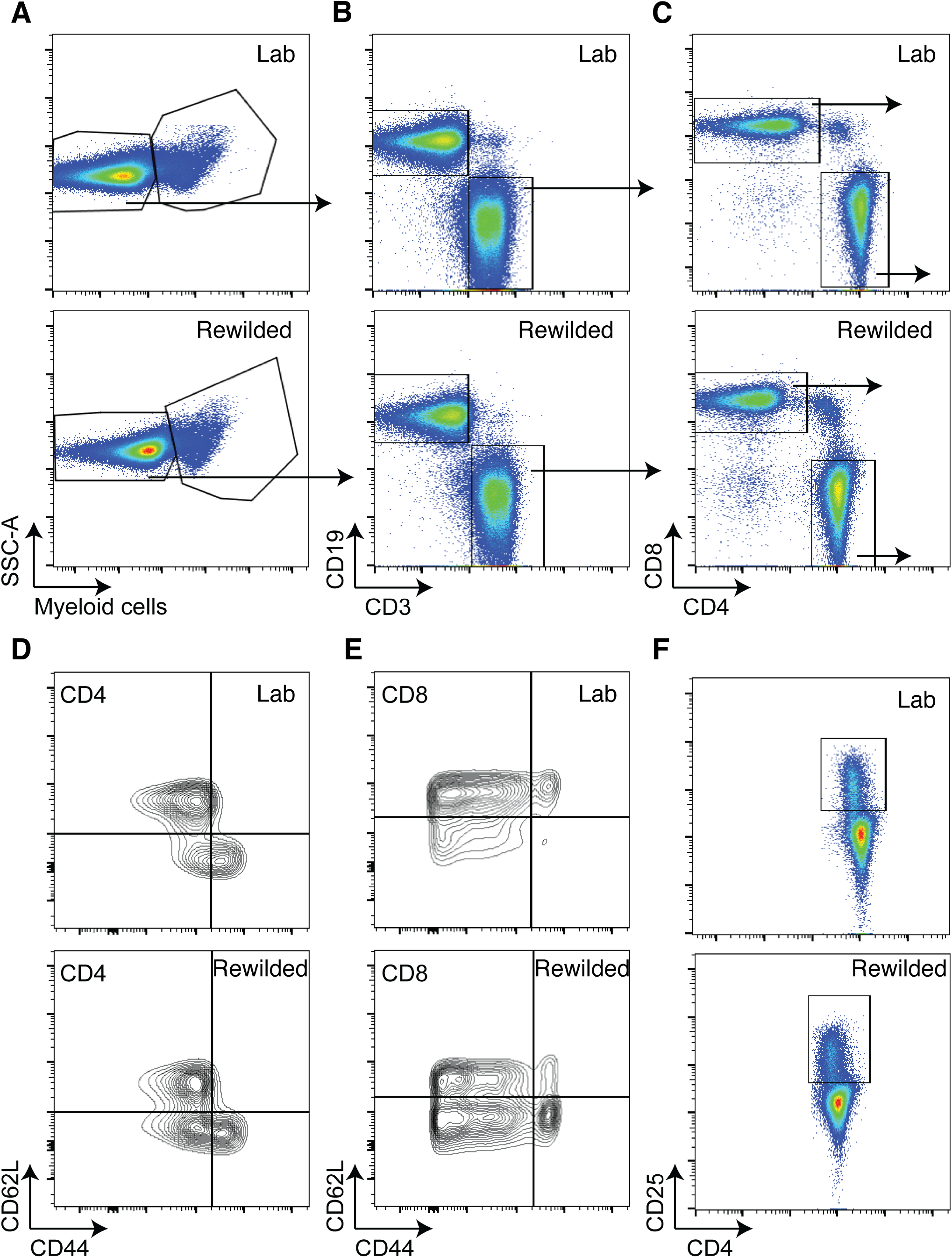
Memory CD4+ and CD8+ T cells are increased in the mesenteric lymph nodes (MLNs) of rewilded mice. (**A** to **F**) Representative flow cytometry plots for immune cell populations after gating for CD45+ cells in the MLNs of lab or rewilded C57BL/6 mice; for the total abundance of myeloid cells (A), B and T cells (B), CD4 and CD8 T cells (C), CD44^lo^CD62L^hi^, CD44^hi^CD62L^hi^, CD44^hi^CD62L^lo^, CD44^lo^CD62L^lo^ CD4 T cells (D), CD44^lo^CD62L^hi^, CD44^hi^CD62L^hi^, CD44^hi^CD62L^lo^, CD44^lo^CD62L^lo^ CD8 T cells (E), and CD25+CD4+ T cells (F).

**Figure S5:**
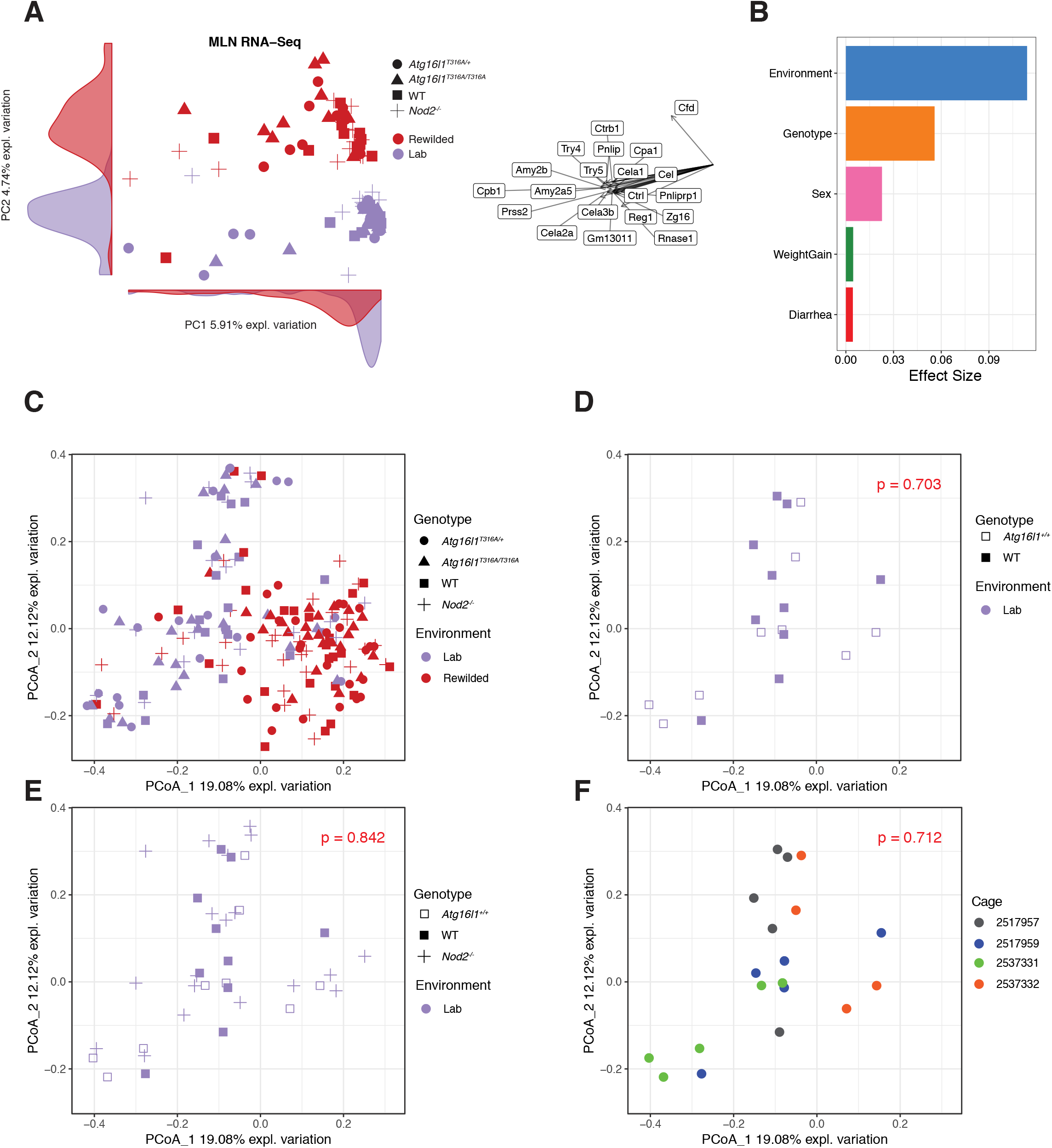
Environment has the greatest effect on variation of transcriptional profiles and microbial composition. (**A** to **B**) Transcriptional profiles of MLNs were determined by Celseq. (**A**) PCA based on gene profiles in the MLN and the density of each population along the principal components (PC). Right panel indicates biplots of the genes projected onto PC1 and PC2. (**B**) Bar plot showing the pseudo R^2^ measure of effect size on the entire distance matrix used to calculate the PCA of gene expression levels. (**C**) PCoA based on Bray Curtis distances of 16s rRNA sequencing profiles in all lab and rewilded mice across all genotypes, lab wild type (WT) and *Atg16l*^+/+^ only, (**E**) lab wild type (WT), *Atg16l*^+/+^ and *Nod2*^−/−^ and (**F**) lab wild type (WT), *Atg16l*^+/+^ and *Nod2*^−/−^ by respective cages. P-values were determined by an Adonis test from Bray Curtis distances.

**Figure S6:**
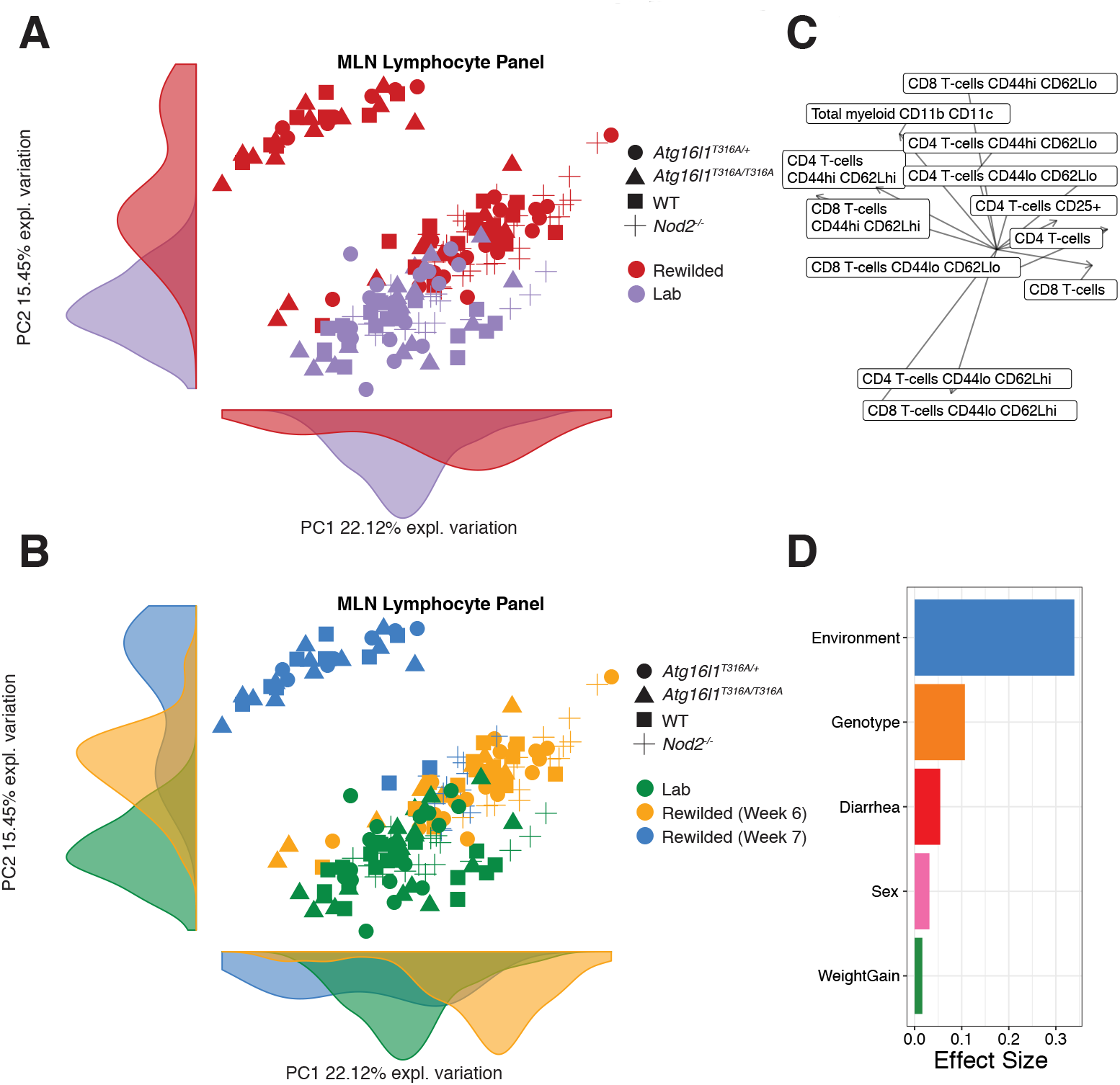
Unmeasured environmental effects drive additional variation of MLN immune cell populations for the rewilded mice. (**A** and **B**) PCA based on lymphoid populations in the MLN and the density of each population along the principal components (PC). In plot (A), dot color coded by rewilded (red) and lab (purple) and in plot (B), dot color coded by lab (green), rewilded week 6 (yellow), rewilded week 7 (blue) (**C**) Biplots of the MLN lymphocyte populations are projected onto PC1 and PC2. (**D**) Bar plot showing the pseudo R^2^ measure of effect size on the entire distance matrix used to calculate the PCA of immune populations.

**Figure S7:**
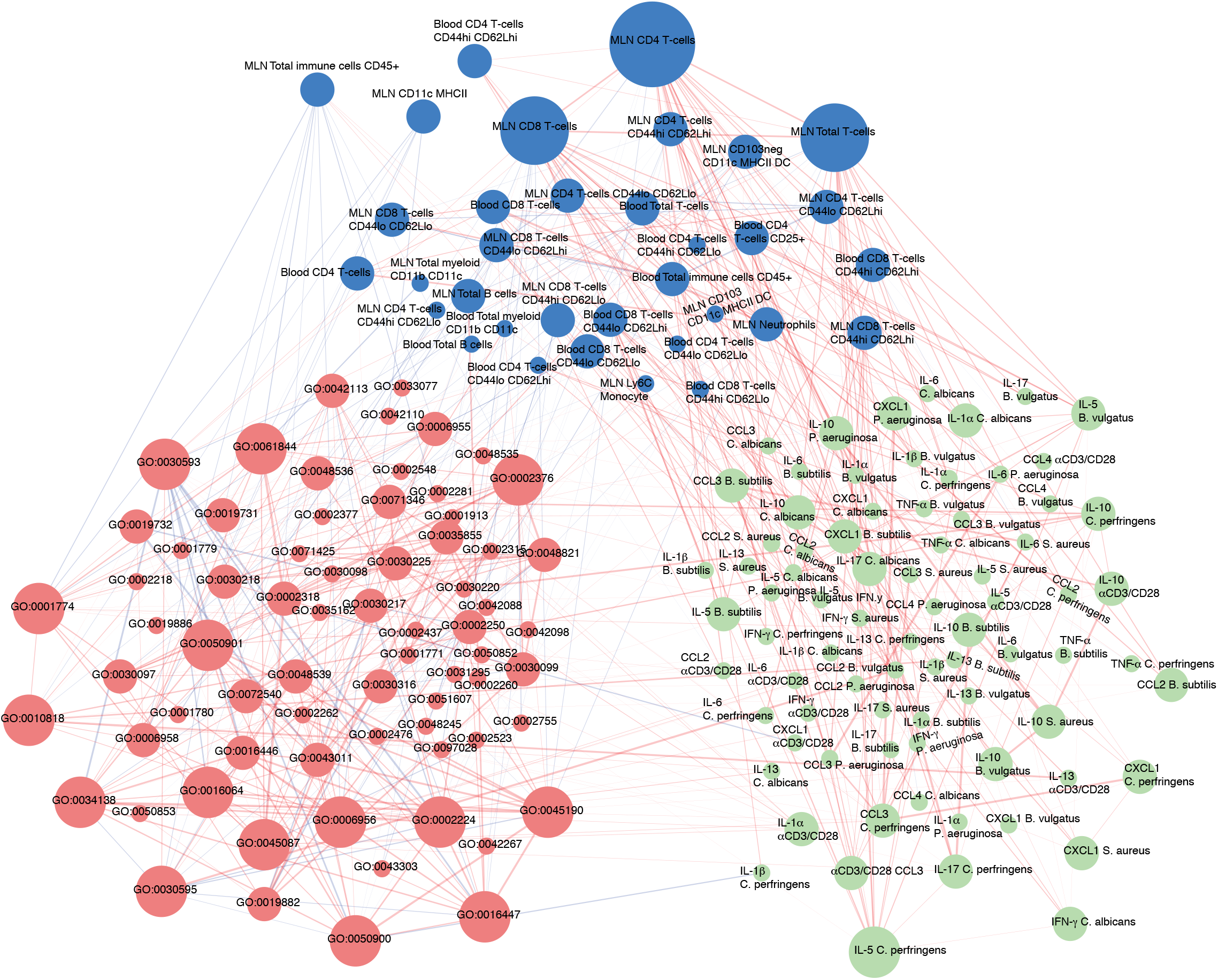
An unsupervised PLS-regression multi-component network model for multi-omic integration. A fully annotated multi-component network as shown in Figure 5A.

