## Supplementary material for "Housing laboratory mice deficient for *Nod2* and *Atg16l1* in a natural environment uncovers genetic and environmental contributions to immune variation": TableS1 and STAR METHODS

**Table S1: Lymphocyte panel**

| **Immune population** | **Gating strategy** |
|---|---|
| Total CD45+ cells | Total cells/Single cells/Live+/CD45+ |
| Total myeloid cells | Total cells/Single cells/Live+/CD45+/CD11b+CD11c+ |
| Total B cells | Total cells/Single cells/Live+/CD45+/CD11b-CD11c-/ CD3-CD19+ |
| Total T cells | Total cells/Single cells/Live+/CD45+/CD11b-CD11c-/ CD3+CD19- |
| Total CD4 T cells | Total cells/Single cells/Live+/CD45+/CD11b-CD11c-/ CD3+CD19-/CD4+CD8- |
| Total CD8 T cells | Total cells/Single cells/Live+/CD45+/CD11b-CD11c-/ CD3+CD19-/CD4-CD8+ |
| $CD44^{lo}CD62L^{hi}$ CD4 T cells | Total cells/Single cells/Live+/CD45+/CD11b-CD11c-/ CD3+CD19-/CD4+CD8-/ $CD44^{lo}CD62L^{hi}$ |
| $CD44^{hi}CD62L^{hi}$ CD4 T cells | Total cells/Single cells/Live+/CD45+/CD11b-CD11c-/ CD3+CD19-/CD4+CD8-/ $CD44^{hi}CD62L^{hi}$ |
| $CD44^{hi}CD62L^{lo}$ CD4 T cells | Total cells/Single cells/Live+/CD45+/CD11b-CD11c-/ CD3+CD19-/CD4+CD8-/ $CD44^{hi}CD62L^{lo}$ |
| $CD44^{lo}CD62L^{lo}$ CD4 T cells | Total cells/Single cells/Live+/CD45+/CD11b-CD11c-/ CD3+CD19-/CD4+CD8-/ $CD44^{lo}CD62L^{lo}$ |
| $CD44^{lo}CD62L^{hi}$ CD8 T cells | Total cells/Single cells/Live+/CD45+/CD11b-CD11c-/ CD3+CD19-/CD4-CD8+/ $CD44^{lo}CD62L^{hi}$ |
| $CD44^{hi}CD62L^{hi}$ CD8 T cells | Total cells/Single cells/Live+/CD45+/CD11b-CD11c-/ CD3+CD19-/CD4-CD8+/ $CD44^{hi}CD62L^{hi}$ |
| $CD44^{hi}CD62L^{lo}$ CD8 T cells | Total cells/Single cells/Live+/CD45+/CD11b-CD11c-/ CD3+CD19-/CD4-CD8+/ $CD44^{hi}CD62L^{lo}$ |
| $CD44^{lo}CD62L^{lo}$ CD8 T cells | Total cells/Single cells/Live+/CD45+/CD11b-CD11c-/ CD3+CD19-/CD4-CD8+/ $CD44^{lo}CD62L^{lo}$ |
| Total CD25+CD4+ T cells | Total cells/Single cells/Live+/CD45+/CD11b-CD11c-/ CD3+CD19-/CD4+CD8-/CD25+CD4+ |

### STAR METODS

#### KEY RESOURCES TABLE

| REAGENT or RESOURCE | SOURCE | IDENTIFIER |
|---|---|---|
| Antibodies | | |
| Pacific Blue anti-mouse CD49b (pan-NK) Antibody | Biolegend | CAT#108918; RRID: AB_2265144 |
| Pacific Blue anti-mouse/human CD11b Antibody | Biolegend | CAT#101224; RRID: AB_755986 |
| Pacific Blue anti-mouse CD11c Antibody | Biolegend | CAT#117322; RRID: AB_755988 |
| Brilliant Violet 421 anti-mouse CD183 (CXCR3) Antibody | Biolegend | CAT#126529; RRID: AB_2563100 |
| Brilliant Violet 510 anti-mouse/rat/human CD27 Antibody | Biolegend | CAT#124229; RRID: AB_2565795 |
| Brilliant Violet 605 anti-mouse/human KLRG1 (MAFA) Antibody | Biolegend | CAT#138419; RRID: AB_2563357 |
| Brilliant Violet 785 anti-mouse CD3ε Antibody | Biolegend | CAT#100355 |
| Brilliant Violet 711 anti-mouse CD127 (IL-7Rα) Antibody | Biolegend | CAT#135035; RRID: AB_2564577 |
| PerCP/Cyanine5.5 anti-mouse CD279 (PD-1) Antibody | Biolegend | CAT#109120; RRID: AB_2566641 |
| APC/Cyanine7 anti-mouse CD4 Antibody | Biolegend | CAT#100414; RRID: AB_312699 |
| PE/Dazzle594 anti-mouse CD19 Antibody | Biolegend | CAT#115554; RRID: AB_2564001 |
| Brilliant Violet 650 anti-mouse CD8a Antibody | Biolegend | CAT#100742; RRID: AB_2563056 |
| Alexa Fluor 488 anti-mouse CD43 Activation-Associated Glycoform Antibody | Biolegend | CAT#121210; RRID: AB_528801 |
| APC anti-mouse CD62L Antibody | Biolegend | CAT#104412; RRID: AB_313099 |
| PE anti-mouse/human CD44 Antibody | Biolegend | CAT#103008; RRID: AB_312959 |
| Alexa Fluor 700 anti-mouse CD69 Antibody | Biolegend | CAT#104539; RRID: AB_2566304 |
| BUV395 Rat Anti-Mouse CD45 Antibody | BD Bioscience | CAT#564279; RRID: AB_2651134 |
| CD25 Monoclonal Antibody (PC61.5), PE-Cyanine7 | eBioscience | CAT#25-0251-82; RRID: AB_469608 |
| Pacific Blue anti-mouse/human CD45R/B220 Antibody | Biolegend | CAT#103227; RRID: AB_2566304 |
| Brilliant Violet 510 anti-mouse CD86 Antibody | Biolegend | CAT#105040; RRID: AB_2315766 |
| Brilliant Violet 605 anti-mouse CD3 Antibody | Biolegend | CAT#100237; RRID: AB_2562039 |
| Brilliant Violet 785 anti-mouse CD69 Antibody | Biolegend | CAT#104543; RRID: AB_2629640 |
| Alexa Fluor 488 anti-mouse CD40 Antibody | Biolegend | CAT#102910; RRID: AB_492852 |
| PerCP/Cyanine5.5 anti-mouse Ly-6G Antibody | Biolegend | CAT#127616; RRID: AB_1877271 |
| APC anti-mouse CD274 (B7-DC, PDL2) Antibody | Biolegend | CAT#127210; RRID: AB_2566345 |

| | | |
|---|---|---|
| APC/Cyanine7 anti-mouse IA/IE Antibody | Biolegend | CAT#107628; RRID: AB_2069377 |
| PE anti-mouse CD274 (B7-H1, PD-L1) Antibody | Biolegend | CAT#124308; RRID: AB_2073556 |
| PE/Dazzle594 anti-mouse CD64 (FCγRI) Antibody | Biolegend | CAT#139320; RRID: AB_2566559 |
| Alexa Fluor 700 anti-mouse F4/80 Antibody | Biolegend | CAT#123130; RRID: AB_2293450 |
| Brilliant Violet 650 anti-mouse CD11c Antibody | Biolegend | CAT#117339; RRID: AB_2562414 |
| BV421 Rat Anti-Mouse Siglec-F Antibody | BD Bioscience | CAT#562681; RRID: AB_2722581 |
| BV711 Rat Anti-Mouse CD103 Antibody | BD Bioscience | CAT#564320; RRID: AB_2738743 |
| PE-Cy7 Rat Anti-Mouse Ly-6C Antibody | BD Bioscience | CAT#560593; RRID: AB_1727557 |
| BUV395 Rat Anti-CD11b Antibody | BD Bioscience | CAT#563553; RRID: AB_2738276 |
| Bacterial and Virus Strains | | |
| *Staphylococcus aureus* | K. Maurer *et al.,* 2015 | USA300 |
| *Pseudomonas aeruginosa* (PAO1) | D. Srivastava *et al.*, 2018 | PAO1 |
| *Bacillus subtilis* | ATCC | ATCC 6633 |
| *Clostridium perfringens* | NCNC | NCTC 10240 |
| *Bacteroides vulgatus* | ATCC | ATCC 8482 |
| *Candida albicans* | Dr. Stefan Feske, NYU | UC820 |
| Chemicals, Peptides, and Recombinant Proteins | | |
| RBC lysis buffer | SANTA CRUZ | Cat#sc-296258 |
| HBSS | Gibco | Cat#14175-095 |
| BSA | Sigma | Cat#A7030 |
| EDTA | Invitrogen | Cat#15575-038 |
| HEPES | Corning | Cat#25-060-CI |
| Sodium pyruvate | Corning | Cat#25-000-CI |
| Critical Commercial Assays | | |
| Live/Dead Fixable Dead Cell Stain Kits | Invitrogen | Cat#L23105 |
| Custom mouse LEGENDplex assay | Biolegend | |
| NucleoSpin Soil Kit | Macherey-Nagel | Cat#740780.250 |
| RNeasy Plus Mini Kit | QIAGEN | Cat#74136 |
| Deposited Data | | |
| 16*S*, ITS, and RNA sequencing reads | NCBI Sequence Read Archive | PRJNA559026 |
| RNA expression counts | Gene Expression Omnibus | GSE135472 |

| | | |
|---|---|---|
| Cytokine and flow cytometry profiles | Github | https://github.com/ruggleslab/RewildedMice |
| Experimental Models: Organisms/Strains | | |
| Mouse: C57BL/6J | The Jackson Laboratory | JAX: 000664 |
| Mouse: *Nod2*$^{-/-}$ | Ramanan et. al., 2014 | |
| Mouse: *Atg16l1*$^{T316A/+}$ | Y. Matsuzawa-Ishimoto *et al.*, 2017 | |
| Mouse: *Atg16l1*$^{T316A/T316A}$ | Y. Matsuzawa-Ishimoto *et al.*, 2017 | |
| Software and Algorithms | | |
| Flowjo 10.4.2 | Flowjo, LLC | https://www.flowjo.com/ |
| Illustrator CC | Adobe | https://www.adobe.com/products/illustrator.html |
| Algorithms: t-SNE | fitsne v1.0.1 package | https://arxiv.org/abs/1712.09005 |
| Algorithms: UMAP | umap-learn v0.3.7 package | https://arxiv.org/abs/1802.03426 |
| Software: Python v3.6.5 | Python.org | https://www.python.org/downloads/release/python-365/ |
| Software: R v3.4.1 | | https://www.r-statistics.com/ |
| Algorithms: principal component analysis | ape v5.2 package | https://www.springer.com/gp/book/9781461417422 |
| Algorithms: Effect size measures | MDMR v0.5.1 package | https://link.springer.com/article/10.1007/s11336-016-9527-8 |
| Algorithms: random forest model | caret v6.0-80 package | R v3.4.1 |
| QIIME2 | Github | https://doc.qiime2.org |

#### LEAD CONTACT AND MATERIALS AVAILABILITY

#### EXPERIMENTAL MODEL AND SUBJECT DETAILS

**Mice and wild enclosure.**

All mouse lines were bred onsite in an MNV/Helicobacter-free specific pathogen free (SPF) facility at NYU School of Medicine to generate littermates from multiple breeding pairs that were randomly assigned to either remain in the institutional vivarium (lab mice) or released into the outdoor enclosures (rewilded mice) to control for the microbiota at the onset of the experiment. *Nod2*$^{-/-}$ and *Atg16l1*$^{T316A/T316A}$ mice on the C57BL/6J background were previously described (Matsuzawa-Ishimoto et al., 2017; Ramanan et al., 2014). *Atg16l1*$^{T316A/T316A}$ mice, *Atg16l1*$^{T316A/+}$, and wild-type (WT) control mice were generated from *Atg16l1*$^{T316A/+}$ breeder pairs, and *Nod2*$^{-/-}$ mice were generated from *Nod2*$^{-/-}$ breeder pairs. Additional C57BL/6J mice were purchased from Jackson Laboratory and bred onsite to supplement WT controls for experiments. 16S microbial diversity at the conclusion of the experiment did not show appreciable differences in microbial composition within the lab populations (Supplementary Figure S5). Outdoor enclosures were previously described (Budischak et al., 2018; Leung et al., 2018) and the protocols for releasing the laboratory mice into the outdoor enclosure facility were approved by Princeton IACUC.

The enclosures consist of replicate outdoor pens, each measuring about 180 $m^2$ and fenced by 1.5-m high, zinced iron walls that are buried >80 cm deep and topped with electrical fencing to keep out terrestrial predators. Aluminum pie plates are strung up to deter aerial predators. A (180 × 140 × 70 cm) straw-filled shed is provided in each enclosure, along with two watering stations and a feeding station, so that the same mouse chow used in the laboratory (PicoLab Rodent Diet 20) was provided ad libitum to all mice. Mice outdoors, however, also had access to food sources found within the enclosures, including berries, seeds, and insects. 26-30 mice of mixed genotypes but the same sex were housed in each enclosure for 6-7 weeks. Longworth traps baited with chow were used to catch mice approximately 2 weeks and 4 weeks after release and again 6-7 weeks after release; for each trapping session, two baited traps were set per mouse per enclosure in the early evening, and all traps were checked within 12 hours. For subsequent microbiome assessment, a fresh stool sample was collected directly from the caught mice, flash frozen on dry ice, and stored at −80℃ until further analysis. Mice were weighed with a spring balance.

30 WT, 29 *Nod2*$^{-/-}$, 31 *Atg16l1*$^{T316A/+}$, and 26 *Atg16l1*$^{T316A/T316A}$ laboratory mice (Total=116) were released into the outdoor enclosure. 19 WT, 19 *Nod2*$^{-/-}$, 20 *Atg16l1*$^{T316A/+}$, 22 *Atg16l1*$^{T316A/T316A}$ matched littermates (Total=80) were maintained in the institutional vivarium for comparison. For rewilded mice, traps were set regularly until the remaining mice were caught and were sampled for fecal microbiota. 25 WT, 28 *Nod2*$^{-/-}$, 27 *Atg16l1*$^{T316A/+}$, and 24 *Atg16l1*$^{T316A/T316A}$ rewilded mice (Total=104) were caught in the final trapping for terminal analyses. All lab control mice were recovered. Euthanasia was performed by CO2 asphyxiation, and blood, MLNs, and intestinal tissue were harvested. Two *Atg16l1*$^{T316A/+}$ rewilded mice failed quality control and were not included in downstream analyses. One *Atg16l1*$^{T316A/+}$ lab mouse was not appropriately processed and excluded in the final meta data table (Table S2; N=79 in lab mice and N=102 in rewilded mice).

#### METHOD DETAILS

**Flow cytometry analysis.**

At harvesting, MLNs were removed and the single-cell suspensions were prepared in FACS buffer (HBSS containing 1% BSA, 1mM EDTA, 20mM HEPES, and 1mM sodium pyruvate). The whole blood were also collected in a heparin containing tube and after centrifuging at 2000 rpm for 5 minutes, the designated plasma from supernatant was removed and stored at −80℃ until all samples were collected and analyzed together. After two rounds of red blood cell lysis with 1x RBC lysis buffer for 5 minutes and wash with FACS buffer, the single-cell suspensions of whole blood cells were ready for the following staining procedure. MLN and whole blood cells were stained for live/dead with blue reactive dye and cell surface markers were labeled with the following antibody panels: Lymphoid panel: CD49b Pacific Blue, CD11b Pacific Blue, CD11c Pacific Blue, CXCR3 Brilliant Violet 421, CD27 Brilliant Violet 510, KLRG1 Brilliant Violet 605, CD3 Brilliant Violet 786, CD127 Brilliant Violet 711, PD1 PerCP/Cy5.5, CD4 APC/Cy7, CD19 PE/Dazzle594, CD8 Brilliant Violet 650, CD43 Alexa Fluor 488, CD62L APC, CD44 PE, CD69 Alexa Fluor 700, CD45 Buv395, CD25 PE/Cy7. Myeloid panel: B220 Pacific Blue, CD86 Brilliant Violet 510, CD3 Brilliant Violet 605,CD69 Brilliant Violet 786, CD40 Alexa Fluor 488, Ly6G PerCP/Cy5.5, PDL2 APC, IA/IE APC/Cy7, PDL1 PE, CD64 PE/Dazzle594, F4/80 Alexa Fluor 700, CD11c Brilliant Violet 650, Siglec-F Brilliant Violet 421, CD103 Brilliant Violet 711, Ly6C PE/Cy7, CD11b Buv395. FACS analyses were performed in a ZE5 cell analyzer (BIO-RAD) and recorded FACS data were analyzed by Flowjo v10.4.2.

**MLN cell stimulation and cytokine profiling**

Single cell suspension of MLN cells were reconstituted in RPMI at $2 \times 10^6$ cells/mL, and 0.1 mL was cultured in 96-well microtiter plates that contained $10^7$ cfu/mL UV-killed microbes, $10^5$ αCD3/CD28 beads, or PBS control. Overnight microbial cultures were reconstituted at $10^8$ cfu/mL prior to irradiation. The stimulated microbes are as following: *Staphylococcus aureus* (Maurer et al., 2015), *Pseudomonas aeruginosa* (PAO1) (kindly provided by Dr. Andrew Darwin, NYU) (Srivastava et al., 2018), *Bacillus subtilis* (ATCC 6633), *Clostridium perfringens* (NCTC 10240), *Bacteroides vulgatus* (ATCC 8482), and *Candida albicans* (UC820, kindly provided by Dr. Stefan Feske, NYU). Supernatants were collected after 2 days and stored at −80°C. Concentrations of IL-1α, IL-1β, IL-4, IL-5, IL-6, IL-10, IL-13, IL-17A, CCL2, CCL3, CCL4, CXCL1, IFN-γ, and TNF-α in supernatants were measured using a custom mouse LEGENDplex assay (Biolegend) according to the manufacturer’s instructions. Plasma concentrations of IL-1α, IL-1β, IL-6, IL-10, RANTES, CCL2, CCL3, CCL4, CCL20, CXCL1, CXCL10, TNFα, GM-CSF were measured using a second custom mouse LEGENDplex assay (Biolegend).

**16S library preparation and sequencing.**

DNA was isolated from stool samples using the NucleoSpin Soil Kit (Macherey-Nagel). Bacterial 16S rRNA gene was amplified at the V4 region using primer pairs and paired-end amplicon sequencing was performed on the Illumina MiSeq system as previously described (Neil et al., 2019). Sequencing reads were processed using the DADA2 pipeline in the QIIME2 software package. Taxonomic assignment was performed against the Silva v132 database. Differential abundance taxa was identified using discrete false-discovery rate (DS-FDR) methodology in different biological groups at a threshold DS-FDR score of 30 (Jiang et al., 2017).

**MLN cell RNA preparation and sequencing**

Frozen samples of single cell suspensions from MLN of lab or rewilded mice were thaw to isolate RNA from approximately $10^6$ cells by RNeasy Plus Mini Kit according to manufacturer's instructions. CEL-seq2 were performed to do RNA sequencing on samples with good RNA qualities (RNA integrity number ≧ 5)

#### QUANTIFICATION AND STATISTICAL ANALYSIS

##### FACS data visualizations by t-distributed stochastic neighbor embedding (t-SNE) and uniform manifold approximation and projection (UMAP)

Flowjo v10.4.2 software and plugin were installed per manufacturer's instructions. The DownSample function in plugin was applied to filter out 1,000 cells on total gated CD45+ cells in blood lymphoid panel staining and the raw channel values from each staining marker were exported to make a FACS value matrix (16 X 1,000) per mouse. FACS data from 2 mice were not passing the data filtrations and 180 FACS value matrixes were concatenated into a giant matrix to perform the downstream t-SNE (fitsne v1.0.1 package (https://arxiv.org/abs/1712.09005), Python v3.6.5) or UMAP (umap-learn v0.3.7 package (https://arxiv.org/abs/1802.03426), Python v3.6.5analysis for the immune cell cluster visualizations. The thresholds were set up for each marker base on expressing distributions across total cells.

**Principal component analysis and effect size measures.** In all cases principal component analysis was performed with the ape (https://www.springer.com/gp/book/9781461417422) v5.2 package in R v3.4.1. Euclidean distances were calculated using base R functions and the subsequent distance matrix was used to determine principle components (PC). Biplots were constructed by projecting the weighted averages of each input feature (immune cell population, cytokine level etc.,) along PC1 and PC2 derived from the biplot.pcoa function from the ape package. Effect size measures were determined using the MDMR v0.5.1 (https://link.springer.com/article/10.1007/s11336-016-9527-8) package in R.

**Per mouse normalization of cytokine production levels.** For each mouse profiled for cytokine production in response to microbial stimuli a PBS control was also sampled. In order to normalize per mouse cytokine production, we calculated the fold change of each cytokine measure to the PBS control. There was no overall difference in baseline cytokine production for any cytokine in response to PBS between lab and rewilded mice.

**Machine learning models for environment and genotype classification.** For multi-omic classification modeling gene expression values, cytokine production levels, immune cell populations and operational taxonomic unit (OTU) counts were normalized by log2-transformation and concatenated together. From ~15,000 genes only the 200 most variable genes, as measured by total variance across all samples, were utilized for modeling. 576 features (200 genes, 104 cytokine measures, 36 immune cell populations and 236 OTUs) from 40 lab and 41 rewilded mice with around 20 mice from each of the four genotypes were supplied to the models. Two random forest models were trained with caret v6.0-80 (R v3.4.1) (5) on a 75% split (61 of 81 mice) with 5-fold cross validation repeated 10 times. The first model was built to classify environment and the second to classify genotype. Both models were evaluated by area under the receiver operator curve (AUC) on the remaining 25% split (20 of 81 mice) left out from the training process. Due to data limitations we did not build an additional model to classify both environment and genotype simultaneously. Feature importance was assessed by the built-in variable importance function varImp within caret.

**Unsupervised multi-omic network model built by sPLS.** To further assess multi-omic relationships between data features (cytokines, immune cell populations, genes and OTUs) an unsupervised network model was constructed. The same 81 mice used to build the classification models were also used to generate this network. The same four data types used to build the classification models were also used with the exception of the 200 most variable genes. Instead we queried the *Mus musculus* Gene Ontology for biological processes related to “immune system process”. From all subsequent child terms, we generated a list of all genes relevant to these processes. Using our gene expression data, we were able to match 954 genes and generate 91 immune system specific gene ontologies. The average expression of all the genes in each ontology were averaged together and used as input into our multi-component sPLS network. In order to generate a multi-omic network of interactions we performed pairwise sparse partial least squares regression as demonstrated by Li et al. (Li et al., 2017) between each of the four data types. After filtering for a covariance threshold of 0.6 in either direction our network consisted of 188 nodes and 577 edges across three data types. None of the OTUs passed this threshold to be included in the network.

#### DATA AND CODE AVAILABILITY

Raw sequence data from 16*S*, ITS, and RNA sequencing experiments are deposited in the NCBI Sequence Read Archive under BioProject accession number PRJNA559026 and gene expression omnibus (GEO) accession number GSE135472. All processing was performed in R and analysis scripts can be found on Github at https://github.com/ruggleslab/RewildedMice

#### METHOD REFERENCE

**Budischak, S.A., Hansen, C.B., Caudron, Q., Garnier, R., Kartzinel, T.R., Pelczer, I., Cressler, C.E., van Leeuwen, A., and Graham, A.L. (2018). Feeding Immunity: Physiological and Behavioral Responses to Infection and Resource Limitation. Front Immunol. 8, 1914. Published online 2018/01/24 DOI: 10.3389/fimmu.2017.01914.**

**Jiang, L., Amir, A., Morton, J.T., Heller, R., Arias-Castro, E., and Knight, R. (2017). Discrete False-Discovery Rate Improves Identification of Differentially Abundant Microbes. mSystems. 2(6). Published online 2017/11/29 DOI: 10.1128/mSystems.00092-17**
**mSystems00092-17 [pii].**

**Leung, J.M., Budischak, S.A., Chung The, H., Hansen, C., Bowcutt, R., Neill, R., Shellman, M., Loke, P., and Graham, A.L. (2018). Rapid environmental effects on gut nematode susceptibility in rewilded mice. PLoS Biol. 16(3), e2004108. Published online 2018/03/09 DOI: 10.1371/journal.pbio.2004108**
**pbio.2004108 [pii].**

**Li, S., Sullivan, N.L., Rouphael, N., Yu, T., Banton, S., Maddur, M.S., McCausland, M., Chiu, C., Canniff, J., Dubey, S., et al. (2017). Metabolic Phenotypes of Response to Vaccination in Humans. Cell. 169(5), 862-877 e817. Published online 2017/05/16 DOI: S0092-8674(17)30477-4 [pii]**
**10.1016/j.cell.2017.04.026.**

**Matsuzawa-Ishimoto, Y., Shono, Y., Gomez, L.E., Hubbard-Lucey, V.M., Cammer, M., Neil, J., Dewan, M.Z., Lieberman, S.R., Lazrak, A., Marinis, J.M., et al. (2017). Autophagy protein ATG16L1 prevents necroptosis in the intestinal epithelium. J Exp Med. 214(12), 3687-3705. Published online 2017/11/02 DOI: jem.20170558 [pii]**
**10.1084/jem.20170558.**
**Maurer, K., Reyes-Robles, T., Alonzo, F., 3rd, Durbin, J., Torres, V.J., and Cadwell, K. (2015). Autophagy mediates tolerance to Staphylococcus aureus alpha-toxin. Cell Host Microbe. 17(4), 429-440. Published online 2015/03/31 DOI: S1931-3128(15)00116-X [pii]**
**10.1016/j.chom.2015.03.001.**

**Neil, J.A., Matsuzawa-Ishimoto, Y., Kernbauer-Holzl, E., Schuster, S.L., Sota, S., Venzon, M., Dallari, S., Galvao Neto, A., Hine, A., Hudesman, D., et al. (2019). IFN-I and IL-22 mediate protective effects of intestinal viral infection. Nat Microbiol. Published online 2019/06/12 DOI: 10.1038/s41564-019-0470-110.1038/s41564-019-0470-1 [pii].**

**Ramanan, D., Tang, M.S., Bowcutt, R., Loke, P., and Cadwell, K. (2014). Bacterial sensor Nod2 prevents inflammation of the small intestine by restricting the expansion of the commensal Bacteroides vulgatus. Immunity. 41(2), 311-324. Published online 2014/08/05 DOI: S1074-7613(14)00241-6 [pii]**
**10.1016/j.immuni.2014.06.015.**

**Srivastava, D., Seo, J., Rimal, B., Kim, S.J., Zhen, S., and Darwin, A.J. (2018). A Proteolytic Complex Targets Multiple Cell Wall Hydrolases in Pseudomonas aeruginosa. MBio. 9(4). Published online 2018/07/19 DOI: mBio.00972-18 [pii]**
**10.1128/mBio.00972-18.**
